# *Lactobacillus johnsonii* Limits Enteropathogenic *Eschericia coli* and *Citrobacter rodentium* Through Biofilm Disruption, Nutrient Competition, and Antimicrobial Metabolites

**DOI:** 10.1101/2025.10.16.682759

**Authors:** Sai Madhuri Vasamsetti, Yasaswi Khaderbad, Novelina Sarmah, Hari Naga Papa Rao Atham, Venkata Ramana Chintalapati, Pavan Kumar Pondugala, Vijay Morampudi

## Abstract

Enteropathogenic *Escherichia coli* is a major cause of childhood diarrhea in underprivileged regions, and its increasing antibiotic resistance underscores the need for non-antibiotic interventions. This study evaluates *Lactobacillus johnsonii* as a probiotic candidate to counter enteropathogenic *Escherichia coli* and its murine surrogate, *Citrobacter rodentium*. *Lactobacillus johnsonii* exhibited robust gastrointestinal resilience, tolerating strong acidity and bile salts (0.3%), and showed enhanced adhesion to human intestinal epithelial cells. Across in vitro assays, live *Lactobacillus johnsonii* inhibited pathogen growth in agar overlay assays more strongly than gentamicin, disrupted biofilms, and displaced adherent enteropathogenic *Escherichia coli* from epithelial surfaces. In antibiotic-treated mice, oral *Lactobacillus johnsonii* reduced *Citrobacter rodentium* burdens in feces, colon, cecum, and spleen by approximately three to four log10 units, mitigated colon shortening, and alleviated histopathological damage, including edema, lymphocyte infiltration, and ulceration. Mechanistic studies revealed complementary modes of action: nutrient competition that reduced pathogen growth by more than fifty percent, and contact-independent killing mediated by secreted, low-molecular-weight factors. Fractionation of cell-free supernatant by fast protein liquid chromatography yielded fractions smaller than seventy-five kilodaltons with potent bactericidal activity; one fraction retained activity for six hours and inhibited enteropathogenic *Escherichia coli* at 30 µg per mL. Untargeted metabolomic profiling of active fractions identified distinct antimicrobial metabolites, including quinine hydrochloride, aloperine, and gamma-glutamylglutamine, alongside chemical classes such as fatty acyls, hydroxy acid derivatives, and carboxylic acids consistent with membrane-disruptive activity. By integrating biofilm disruption, competitive exclusion, and metabolite-mediated killing with demonstrated efficacy in vivo, *Lactobacillus johnsonii* emerges as a promising biotherapeutic for managing diarrheal diseases caused by attaching-and-effacing pathogens and merits further characterization of its active small molecules and translational evaluation.

**Author Summary:** Diarrheal disease still harms millions of children, and growing antibiotic resistance makes treatment harder. We asked whether a friendly gut bacterium, *Lactobacillus johnsonii*, could protect against harmful microbes without relying on antibiotics. First, we tested simple but key questions: can this probiotic survive the harsh journey through the stomach and small intestine, can it attach to the gut lining, and can it push back against disease-causing bacteria?

We found that *Lactobacillus johnsonii* survives strong acid and bile and sticks well to human intestinal cells. In lab dishes, live cells slowed pathogen growth, broke up their protective biofilms (the sticky layers that help microbes persist), and even dislodged bacteria that had already attached to cells. We then moved to a mouse model of diarrheal infection using *Citrobacter rodentium*, which mimics important features of human disease. Giving *Lactobacillus johnsonii* by mouth lowered bacterial counts in the gut and tissues, reduced tissue damage, and improved overall colon health.

Finally, we explored how it works. We found two main actions: the probiotic competes with pathogens for nutrients, and it releases small natural compounds that can directly kill them. Together, these results support *Lactobacillus johnsonii* as a practical, affordable, non-antibiotic option that merits careful testing in people, especially in communities with the greatest burden of diarrheal disease.

**Graphical Abstract:** Schematic overview of the study design and key findings. A high-resolution image and legend are provided.

Mechanism of *Lactobacillus johnsonii*-mediated protection against EPEC and *Citrobacter rodentium* in the gut
**Left (Pathogenic Effects):** Colonization by EPEC or *C. rodentium* disrupts the intestinal mucus barrier, induces attaching and effacing (A/E) lesions on epithelial cells, and triggers pro-inflammatory cytokine release. These events promote biofilm formation, immune cell infiltration (including dendritic cells, macrophages, neutrophils, and T cells), and epithelial ulceration, ultimately exacerbating intestinal inflammation.
**Right (Protective Effects):** *L. johnsonii* counteracts these pathogenic effects through secretion of antimicrobial metabolites, competition for nutrients, and enhanced mucosal adherence. Collectively, these mechanisms inhibit biofilm formation, reduce pathogen burden, and maintain epithelial barrier integrity.

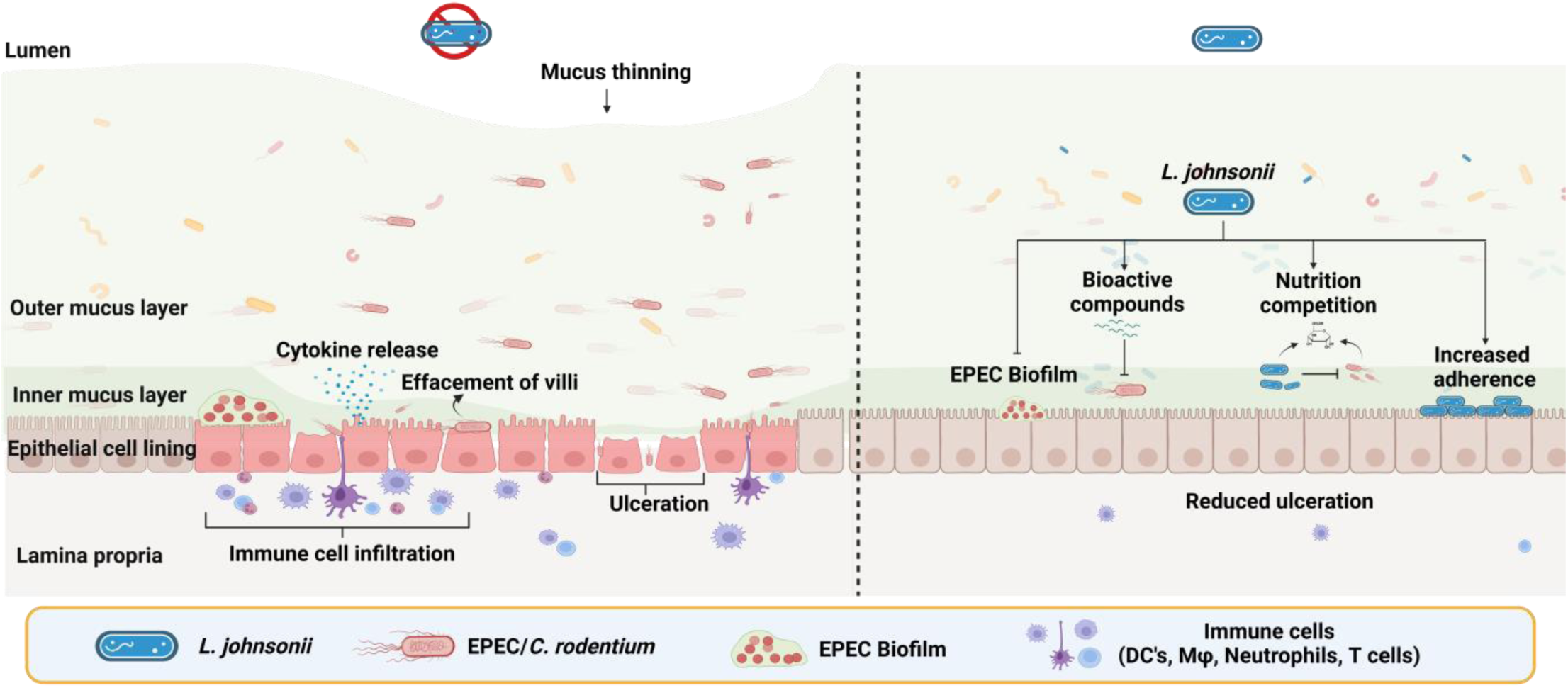

## Introduction

Diarrheal diseases remain a leading cause of morbidity and mortality worldwide, particularly in low- and middle-income countries. The World Health Organization estimates ∼1.7 billion cases annually and more than 525,000 deaths among children under five, making diarrhea a top cause of childhood mortality [1]. South Asia bears a disproportionate burden, accounting for over 50% of global childhood diarrheal cases [2]. In India, diarrheal diseases contribute to ∼100,000 child deaths each year, with enteric bacterial pathogens such as enteropathogenic *Escherichia coli* (EPEC) playing a major role [3].

EPEC is an attaching-and-effacing (A/E) pathogen whose virulence depends on a type III secretion system that translocates bacterial effectors into epithelial cells, producing actin-rich pedestals, disrupting tight junctions, and driving intestinal inflammation [4]. Chronic or recurrent EPEC infection is linked to a three- to fivefold elevated risk of stunting and malnutrition due to impaired absorption and persistent enteropathy [5]. In India, multicenter surveillance has detected EPEC in 12–15% of hospitalized pediatric diarrhea cases [6]. The rise of multidrug-resistant EPEC further complicates treatment, with high resistance reported to commonly used agents such as ampicillin, trimethoprim-sulfamethoxazole, and fluoroquinolones [7]. These trends highlight the need for non-antibiotic approaches..

Probiotics defined as live microorganisms that confer health benefits when administered in adequate amounts offer one such strategy. Reported mechanisms include competitive exclusion of pathogens, strengthening of barrier function, production of antimicrobial compounds, and immunomodulation [8]. Although strains such as *Lactobacillus rhamnosus* GG and *Lactobacillus plantarum (L. plantarum)* can shorten diarrheal duration, benefits are strain-specific, motivating evaluation of additional candidates [9, 10].

*Lactobacillus johnsonii (L. johnsonii)*, a member of the *Lactobacillus delbrueckii* group, has drawn attention for acid and bile tolerance, epithelial adherence, and antimicrobial activity [11]. Genomic and functional studies indicate that some *L. johnsonii* strains produce bacteriocins and other antimicrobial metabolites and express surface-layer proteins that enhance mucosal adhesion [12–15]. Notably, *L. johnsonii* can disrupt pre-formed biofilms via exopolysaccharide-mediated interference with quorum-regulated behaviors, an attribute relevant to persistent infections [16, 17]. Clinical observations further support its safety, including reduced *Helicobacter pylori* colonization in children receiving *L. johnsonii* preparations [18]. At the same time, intrinsic resistance to certain antibiotics (for example, aminoglycosides in some lactobacilli) and susceptibility to others vary by strain [19, 20], underscoring the importance of strain-level characterization.

To address the gap in evidence for A/E pathogens, we investigated the probiotic potential of *L. johnsonii* against EPEC. Because human challenge is not feasible, we used *Citrobacter rodentium (Citrobacter rodentium)*, a murine A/E pathogen that mirrors EPEC in genome content, virulence mechanisms, and disease pathology [21]. *C. rodentium* infection in mice recapitulates key features of EPEC-induced disease, including epithelial hyperplasia, mucosal inflammation, and barrier disruption [22]. We assessed classical probiotic traits (acid/bile tolerance, adhesion, biofilm inhibition, and competitive exclusion) and dissected secreted antimicrobial activities using fast protein liquid chromatography and untargeted metabolomics. Finally, we tested efficacy in antibiotic-perturbed C57BL/6 mice infected with *C. rodentium*, quantifying pathogen burden and colonic pathology. Together, these studies define multifaceted antimicrobial and gut-protective actions of *L. johnsonii* and support its development as a biotherapeutic candidate against multidrug-resistant A/E pathogens.

## Results

### *L. johnsonii* exhibits cysteine-enhanced growth and tolerance to gastrointestinal stressors

To confirm the identity of the probiotic isolate, genomic DNA was first amplified with *L. johnsonii* specific primers, yielding a distinct 126 bp amplicon (Fig 1B). Further validation was performed by 16S rRNA gene sequencing. The sequence exhibited >98% similarity to *L. johnsonii* strain 2317 and has been deposited in GenBank (accession no. PV739486). A schematic of the experimental workflow used to evaluate the strain’s growth profile and physiological robustness is shown in Fig 1A.

**Fig 1.**
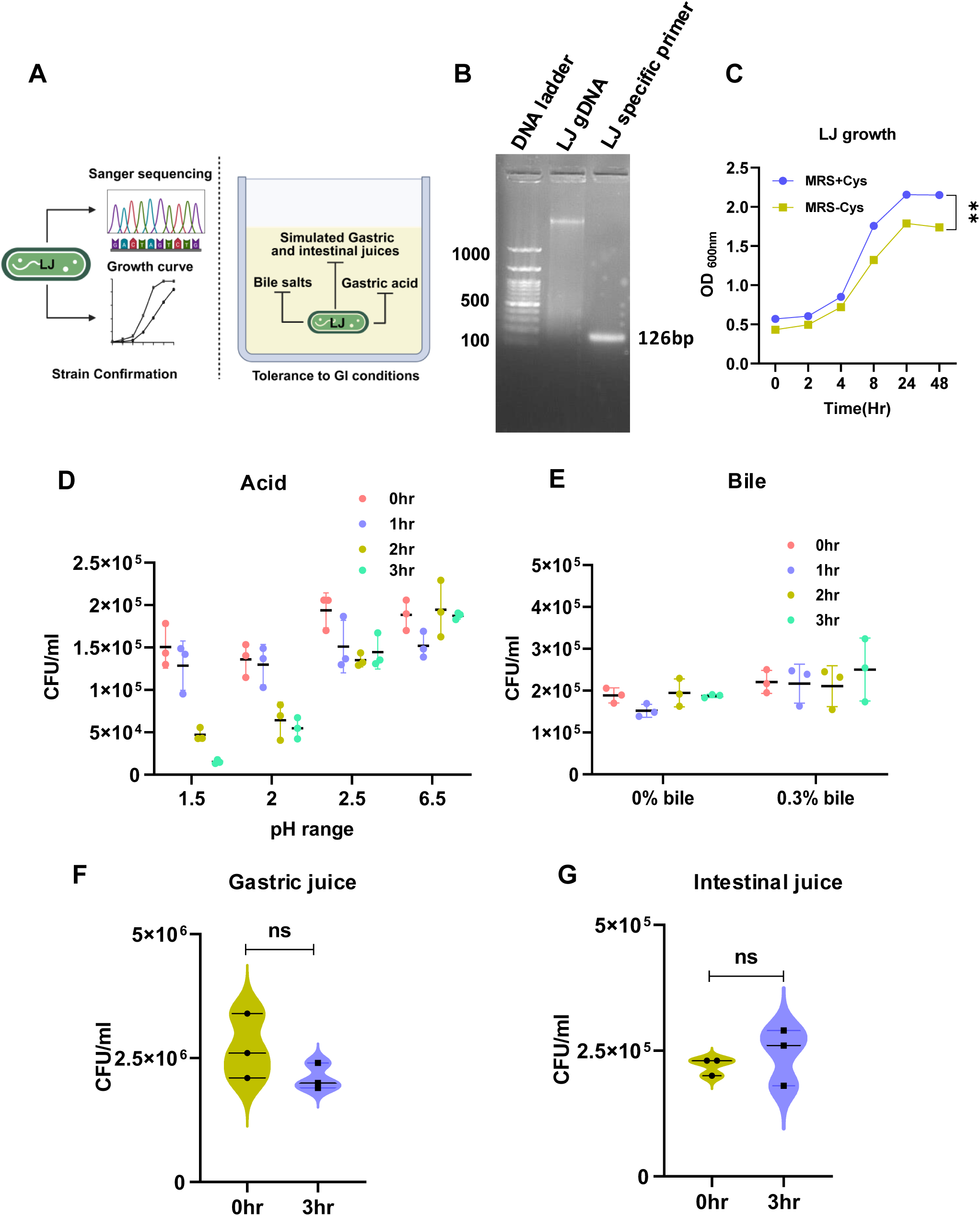
Characterization of *L. johnsonii* for growth, stress resistance, and enzymatic tolerance. (A) Schematic of the experimental workflow for strain confirmation and evaluation of stress tolerance. (B) PCR amplification using *L. johnsonii*-specific primers yields a 100 bp product. Lane 1: 100 bp DNA ladder; lane 2: *L. johnsonii* genomic DNA (template); lane 3: amplification with *L. johnsonii*-specific primers. (C) Growth kinetics in de Man, Rogosa and Sharpe (MRS) broth supplemented with 0.05% (w/v) cysteine (MRS + Cys, blue) or without supplementation (MRS - Cys, yellow). (D) Acid tolerance of *L. johnsonii* assessed at pH 1.5, 2.0, 2.5, and 6.5 over 0-3 h. (E) Viability following exposure to 0.3% (w/v) bile salts for 0-3 h. (F) Survival in simulated gastric juice containing 3.0 g/L pepsin at pH 2.5. (G) Survival in simulated intestinal juice containing 1.0 g/L trypsin and 1.8% (w/v) bile salts at pH 8.0. Data represent mean ± SEM from ≥3 independent biological replicates. Statistical analysis was performed using one-way ANOVA with Dunnett’s multiple comparisons test; \*\**p* < 0.01; ns, not significant.

*L. johnsonii* exhibited significantly enhanced growth in cysteine-supplemented MRS medium (MRS + Cys) compared to the standard formulation (Fig 1C), consistent with observations in *L. plantarum* (S1 Fig). Bacterial CFU counts at OD600 = 1 were quantified for all strains (S1 Table). To assess gastrointestinal stress tolerance, *L. johnsonii* was exposed to acidic conditions (pH 1.5–2.5). The strain remained viable at pH 1.5 and 2.0 for up to 1 hour, and survival extended up to 3 hours at pH 2.5 (Fig 1D). Similarly, viability was maintained after 3-hour exposure to 0.3% bile salts (Fig 1E). When challenged with simulated gastric and intestinal fluids, *L. johnsonii* showed minimal reduction in CFU after 3 hours, indicating strong resistance to pepsin and trypsin (Figs 1F and 1G). Collectively, these results underscore the strain’s ability to survive gut-like conditions and support its potential for in vivo colonization.

### *L. johnsonii* demonstrates antimicrobial and anti-biofilm activity against EPEC

To assess the ability of *L. johnsonii* to inhibit EPEC growth, we conducted an agar overlay assay and a biofilm inhibition assay, with the experimental workflow summarized in Fig 2A. *L. johnsonii* produced a distinct zone of inhibition against EPEC, with a significantly larger diameter compared to gentamicin (30 µg/mL), indicating strong antimicrobial activity (Figs 2B–C). To evaluate anti-biofilm effects, EPEC was co-cultured with live or heat-killed *L. johnsonii* and assessed via crystal violet staining. Only the live strain significantly reduced EPEC biofilm biomass by ∼60% (Fig 2D). Given that ≥50% biofilm inhibition or ≥1 log CFU reduction is considered biologically relevant in probiotic–pathogen interaction studies, this reduction represents a meaningful effect. Correspondingly, viable EPEC counts were markedly lower in the presence of both live and heat-killed *L. johnsonii*, as determined by CFU enumeration (Fig 2E). The partial retention of inhibitory effect with heat-killed cells suggests that both secreted metabolites and structural cell components contribute to EPEC inhibition. In contrast, the biofilm-forming capacity of *L. johnsonii* remained unaffected by EPEC co-culture (Fig 2F). These findings suggest that *L. johnsonii* exerts contact- and secretome-dependent inhibition of EPEC growth and biofilm formation while maintaining its own biofilm integrity.

**Fig 2.**
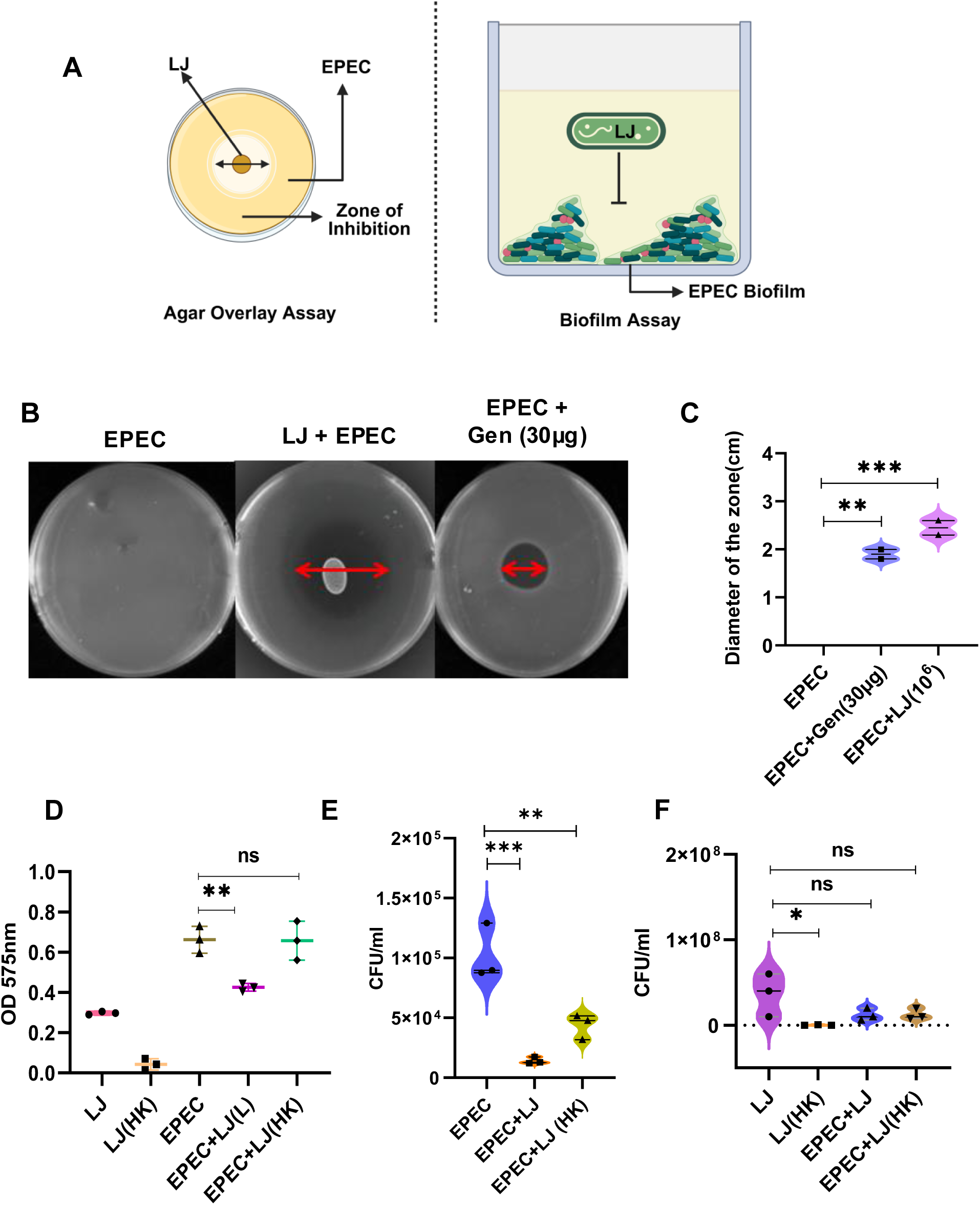
Antimicrobial and anti-biofilm activity of *L. johnsonii* against *EPEC*. (A) Schematic representation of the agar overlay and biofilm inhibition assay workflows. (B) Representative agar overlay plates showing inhibition zones for: (i) EPEC alone, (ii) EPEC overlaid on *L. johnsonii* (LJ), and (iii) EPEC treated with gentamicin (30 µg/mL). (C) Quantification of inhibition zone diameters (cm) for each treatment. (D) Crystal violet assay quantifying EPEC biofilm biomass after 48 h co-culture with live *L. johnsonii* (LJ) or heat-killed *L. johnsonii* (HK-LJ). (E) Enumeration of viable EPEC cells (CFU/mL) recovered from biofilms formed in the presence or absence of LJ or HK-LJ. (F) Enumeration of viable LJ cells (CFU/mL) in biofilms after 48 h co-culture with EPEC. All experiments were performed in triplicate and independently repeated at least three times (n = 3). Data are presented as mean ± SEM. Statistical analysis was performed using one-way ANOVA with Dunnett’s multiple comparisons test; \**p* < 0.05; \*\**p* < 0.01; \*\*\**p* < 0.001; ns- not significant.

### *L. johnsonii* disrupts established *EPEC* colonization but does not prevent initial attachment

To evaluate the functional attributes of *L. johnsonii*, assays were performed to assess its adherence to HCT-116 cells, antibiotic resistance profile, and its ability to inhibit EPEC through exclusion, displacement, and competition mechanisms, as illustrated in Fig 3A. To assess cytotoxicity induced by *L. johnsonii* and EPEC in HCT-116 cells, lactate dehydrogenase (LDH) release was measured at varying multiplicities of infection (MOI). Consistent with previous findings, a significant increase in LDH levels was observed at higher MOIs of *L. johnsonii* and EPEC (S2A and B Figs). An MOI of 25 for both *L. johnsonii* and EPEC was identified as a safe threshold, which was subsequently used for all further experiments. The adherence of EPEC, *L. plantarum*, and *L. johnsonii* to HCT-116 cells was evaluated after 6 hours of incubation at an MOI of 1:25. *L. plantarum*, previously demonstrated to possess antimicrobial activity against EPEC, served as a control. Among the tested strains, *L. johnsonii* exhibited the highest adherence to HCT-116 cells (3.35%) compared to EPEC (1.9%) and *L. plantarum* (1.65%) (Fig 3B). To assess antibiotic susceptibility, *L. johnsonii* and *L. plantarum* were tested against four antibiotics following EUCAST guidelines (S2 Table). Similar to *L. plantarum*, *L. johnsonii* exhibited resistance to kanamycin and gentamicin, while remaining susceptible to ampicillin and vancomycin (Fig 3C).

**Fig 3.**
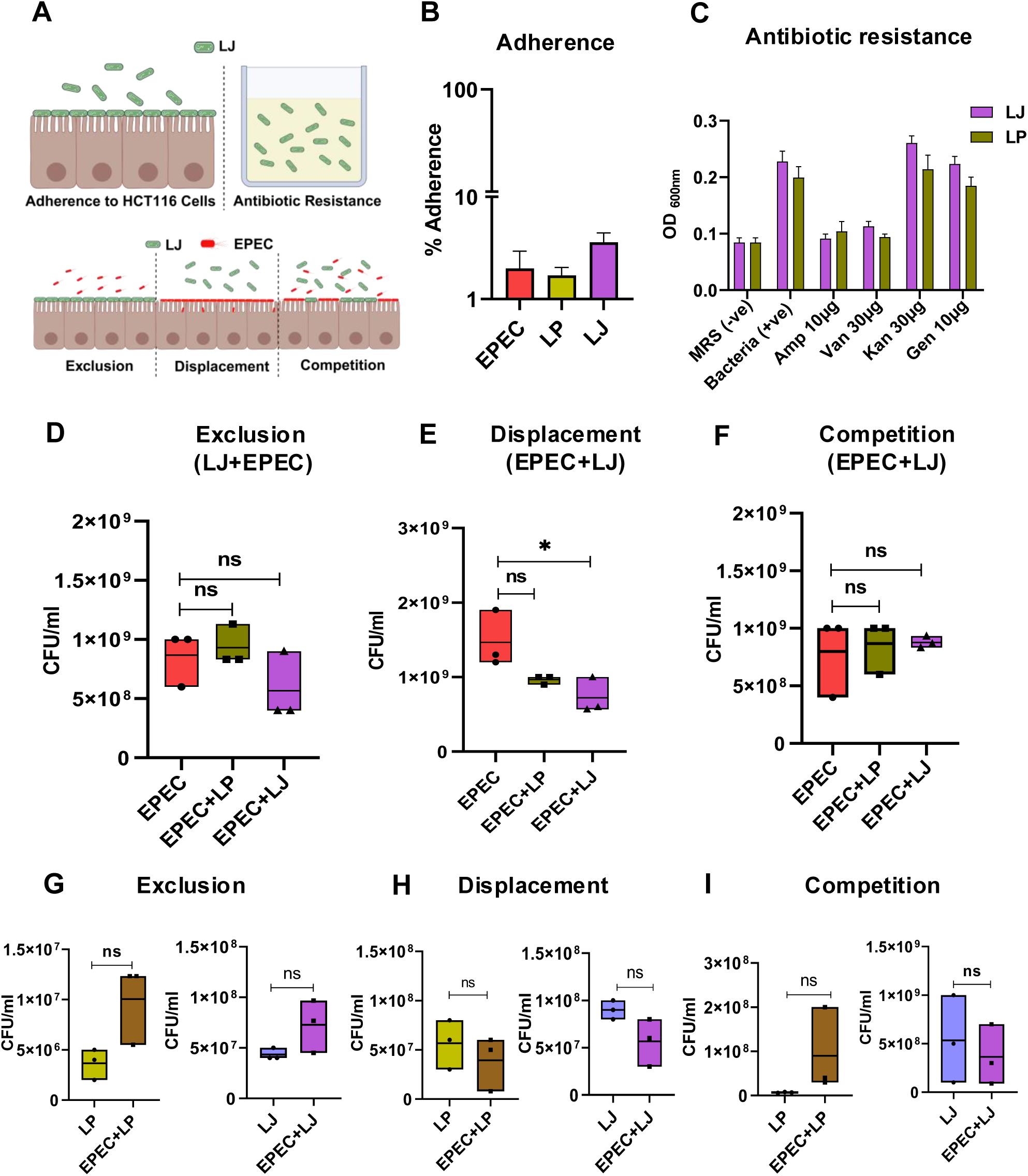
Protective effects of *L. johnsonii* against EPEC attachment and colonization. (A) Schematic representation of experimental assays evaluating the ability of *L. johnsonii* (LJ) to inhibit EPEC adherence and colonization of HCT-116 intestinal epithelial cells. (B) Adherence rates (% of inoculum) of LJ, *Lactobacillus plantarum* (LP), and EPEC to HCT-116 monolayers after 6 h incubation at a multiplicity of infection (MOI) of 1:25. (C) Antibiotic susceptibility profiles of LJ and LP against ampicillin (10 µg), vancomycin (30 µg), kanamycin (30 µg), and gentamicin (10 µg). (D) Exclusion assay: HCT-116 cells pre-incubated with LJ or LP for 3 h, followed by EPEC infection for 3 h. (E) Displacement assay: cells pre-infected with EPEC for 3 h, followed by treatment with LJ or LP for 3 h. (F) Competition assay: LJ or LP co-incubated with EPEC on HCT-116 cells for 6 h. (G-I) Growth of LJ and LP during (G) exclusion, (H) displacement, and (I) competition assays, expressed as CFU/mL. All experiments were performed in triplicate and repeated independently three times (n = 3). Data are presented as mean ± SEM. Statistical analysis was performed using one-way ANOVA followed by Dunnett’s multiple comparisons test;: \**p* < 0.05; ns- not significant.

The protective effects of *L. johnsonii* against EPEC were further evaluated using exclusion, displacement, and competition assays at an MOI of 1:25. In the exclusion assay, pre-treatment with *L. johnsonii* did not significantly reduce EPEC growth, indicating that *L. johnsonii* does not inhibit initial EPEC attachment (Fig 3D). However, in the displacement assay, a significant reduction in EPEC growth was observed following treatment with *L. johnsonii*, suggesting its ability to disrupt established EPEC colonization (Fig 3E). In contrast, the competition assay, in which *L. johnsonii* and EPEC were co-incubated, showed no significant changes in bacterial growth, indicating that *L. johnsonii* does not competitively inhibit EPEC under these conditions (Fig 3F). Furthermore, to determine the impact of EPEC infection on Lactobacillus growth, we analyzed the proliferation of *L. johnsonii* and *L. plantarum* during EPEC infection. Interestingly, co-incubation during the exclusion assay appeared to support the proliferation of both *L. johnsonii* and *L. plantarum*, suggesting mutual tolerance or potential niche adaptation (Figs 3G–I). These findings suggest that while *L. johnsonii* does not prevent EPEC adhesion, it can displace pre-attached EPEC, which may contribute to its protective role against enteropathogenic infections

### Eradication of *L. johnsonii* and reduction in total bacteria following antibiotics treatment

Following the confirmation of *L. johnsonii*’s antimicrobial activity against EPEC in vitro, its effect on *C. rodentium* was assessed in a murine model. Stool samples from naïve C57BL/6 mice were analyzed via PCR using *L. johnsonii* specific primers. Agarose gel electrophoresis of individual and pooled DNA samples revealed distinct *L. johnsonii* bands, confirming its presence in the gut microbiota of untreated mice (Fig 4A). To evaluate the potential of *L. johnsonii* against *C. rodentium*, pre-existing *L. johnsonii* in the mice was eliminated and the gut microbiota composition perturbed by administering a broad-spectrum antibiotic cocktail via a single oral gavage followed by a seven-day course in drinking water (Fig 4B and S3 Table). PCR analysis of stool samples from antibiotic-treated (Abx) mice revealed the absence of *L. johnsonii* DNA bands as early as day 2 (S3A and B Figs.), with complete eradication by day 7 (Fig 4C). In contrast, *L. johnsonii* remained detectable in the antibiotic-untreated control group throughout the experiment. Additionally, real-time PCR using eubacterial primers demonstrated a progressive decline in total bacterial load in the antibiotic-treated group from day 2 to day 7, whereas the control group maintained a stable bacterial population (Fig 4D). These findings indicate that antibiotic treatment effectively eradicates *L. johnsonii* while significantly reducing overall bacterial abundance in the murine gut.

**Fig 4.**
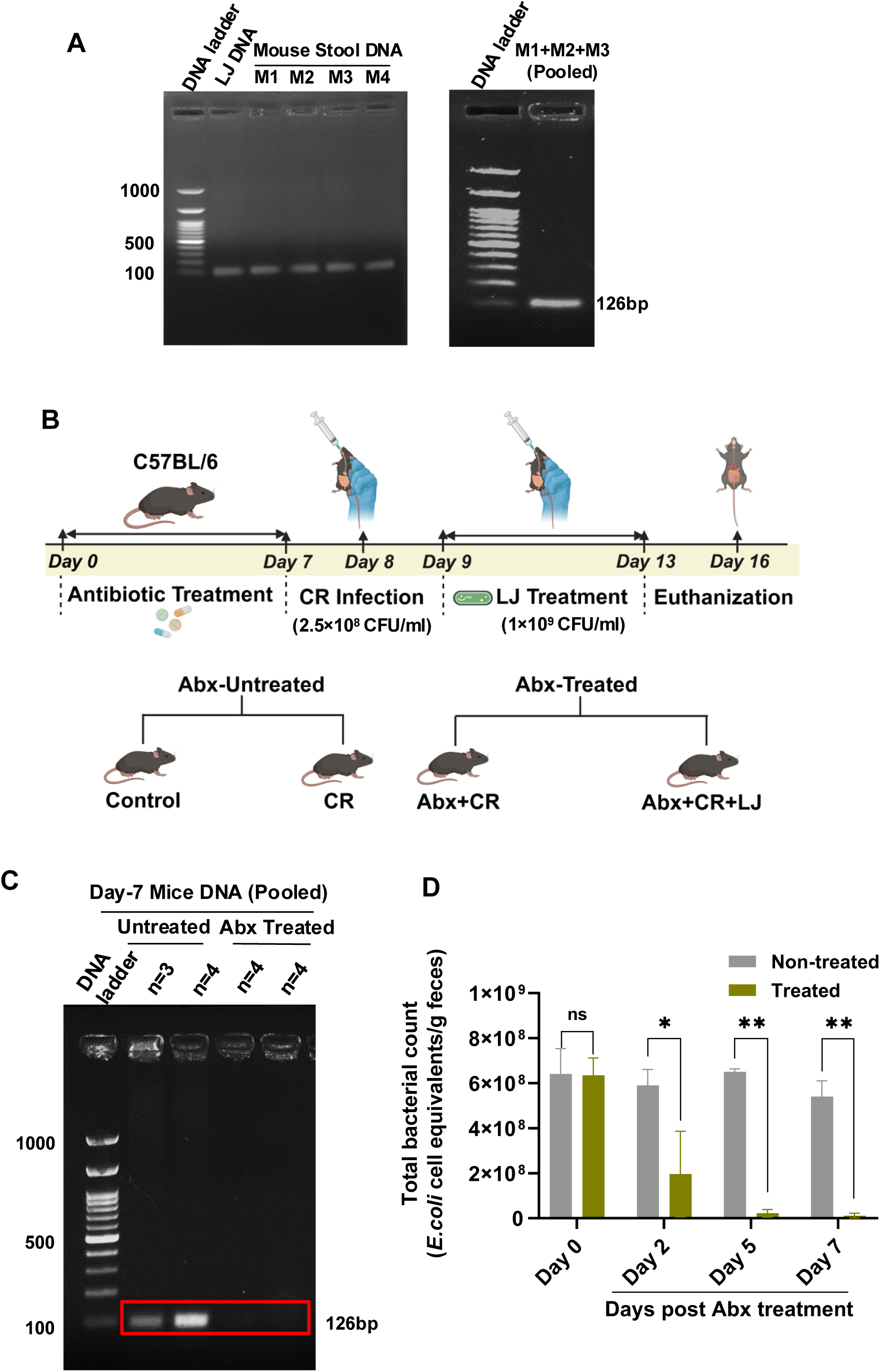
Detection of *L. johnsonii* in naïve mice and its clearance following antibiotic treatment. (A) Agarose gel electrophoresis showing *L. johnsonii*-specific PCR amplification (100 bp amplicon) from fecal DNA of individual naïve mice (M1-M3) and pooled stool DNA. Lane 1: 100 bp DNA ladder; Lane 2: LJ specific primer; lanes 3-6: DNA from individual mice; lane 7: 100 bp DNA ladder and lane 8: pooled DNA from M1-M3. (B) Experimental timeline: mice received a single oral gavage of antibiotic cocktail (Abx) on Day 0, followed by antibiotic-supplemented drinking water (Days 1-7). On Day 8, mice were orally infected with *C. rodentium* (CR). From Days 9-13, the probiotic-treated group received daily LJ (10⁹ CFU in 100 µL PBS). Mice were euthanized on Day 16 (8 days post-infection). Experimental groups: Control (no Abx, no CR), CR (no Abx), Abx+CR, and Abx+CR+LJ. (C) Agarose gel showing complete loss of the *L. johnsonii*-specific 100 bp PCR product by Day 7 in Abx-treated mice. Lane 1: 100 bp ladder; lanes 2-5 correspond to pooled mouse stool DNA samples. (D) Quantitative PCR analysis of total eubacterial 16S rRNA gene copies in fecal DNA from Day 0 to Day 7 post-antibiotic treatment using the standard curve method. Data are mean ± SEM (n = 4 mice/group). Statistical analysis: two-way ANOVA with Dunnett’s multiple comparisons test; \**p* < 0.05; \*\**p* < 0.001; ns- not significant.

### *L. johnsonii* mitigates *C. rodentium* induced colonic pathology

To assess disease severity and therapeutic efficacy, clinical and histological evaluations were conducted in C57BL/6 mice infected with *C. rodentium* (Fig 5A). Body weight changes were monitored during the seven-day antibiotic treatment. Mice receiving antibiotics exhibited noticeable weight loss compared to the untreated control group (Fig 5B). However, upon cessation of antibiotic-containing water, the mice regained weight (Fig 5C). By the end of the treatment period, no significant weight differences were observed, and all four experimental groups maintained a 100% survival rate until the day of sacrifice (Fig 5D). To assess *C. rodentium* induced colonic inflammation, colon length was measured post-sacrifice on day 8 post-infection. Macroscopic evaluation revealed significant colon shortening in *C. rodentium* infected mice compared to naïve controls. Interestingly, *L. johnsonii* treated mice exhibited significantly longer colons than untreated *C. rodentium* infected mice, suggesting a protective effect (Figs 5E–F).

**Fig 5.**
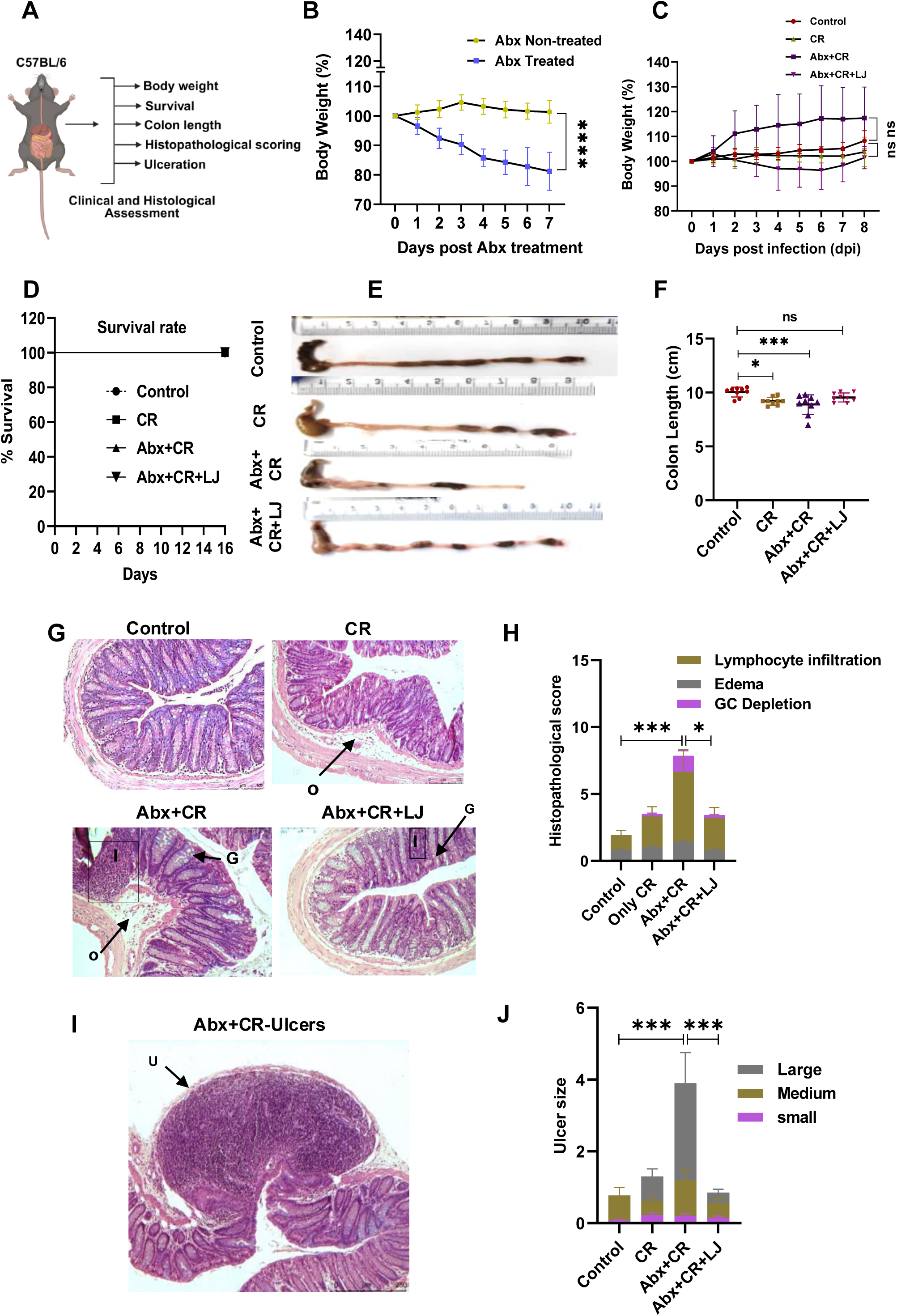
*L. johnsonii* alleviates *Citrobacter rodentium*-associated colonic inflammation. (A) Schematic of clinical and histological assessments used to evaluate disease severity and therapeutic outcomes in C57BL/6 mice. (B) Body weight change (%) during 7 days of antibiotic (Abx) administration in treated versus untreated mice. (C) Body weight recovery following withdrawal of Abx-containing water. (D) Survival percentage of all experimental groups up to the day of sacrifice. (E) Representative macroscopic images of colons at 8 days post-infection (DPI). (F) Quantification of colon length (cm) at 8 DPI; Abx+CR mice exhibited significant shortening compared to controls, with recovery in Abx+CR+LJ mice. (G) Representative hematoxylin and eosin (H&E)-stained colon sections from Control, CR, Abx+CR, and Abx+CR+LJ groups at 8 DPI. (H) Histopathological scoring (maximum score = 20) assessing lymphocyte infiltration, submucosal edema, and goblet cell depletion. (I) Representative H&E-stained image showing extensive ulceration in the Abx+CR group. (J) Quantification of ulcer frequency and size (percentage of cross-sectional tissue affected) in colonic tissue at 8 DPI. Data are mean ± SEM (n = 9-10 mice/group). Statistical analysis was performed using Tukey’s multiple comparisons test; \**p* < 0.05; \*\*\**p* < 0.0001; \*\*\*\**p* < 0.0001; ns- not significant. I- inflamamtory cells, G- goblet cells, O- oedema, and U- ulcer.

Histological analysis on day 8 post-infection demonstrated pronounced colonic inflammation in antibiotic-treated mice with *C. rodentium* infection, which was significantly reduced following daily administration of *L. johnsonii* (Fig 5G). Pathological scoring of colonic tissue further confirmed severe inflammation, characterized by a substantial influx of inflammatory cells, submucosal edema, goblet cell depletion, and disrupted mucosal architecture in infected mice (Fig 5H). Notably, antibiotic-treated mice exhibited more severe ulcerative damage compared to those receiving *L. johnsonii* (Figs 5I–J). Furthermore, immunostaining of colonic sections revealed significantly higher neutrophil infiltration at the mucosal surface in antibiotic-treated, infected mice compared to those receiving probiotic treatment (S4A Fig.). However, Ki-67 staining showed no significant differences in cell proliferation between the two treatment groups (S4B Fig). Overall, the *C. rodentium* infected, antibiotic-treated group exhibited significantly higher pathology scores than control mice. These findings demonstrate that *L. johnsonii* alleviates *C. rodentium* induced colonic pathology and inflammation.

### *L. johnsonii* reduces *C. rodentium* colonization and attenuates antibiotic-exacerbated gut pathology

To evaluate whether *L. johnsonii* can suppress *C. rodentium* colonization and mitigate antibiotic-enhanced pathogen burden, bacterial colony-forming units (CFUs) were quantified in stool and tissue samples (Fig 6A). Fecal analysis revealed that *C. rodentium* CFUs were markedly elevated in antibiotic-treated mice compared to those without antibiotic exposure, indicating enhanced susceptibility following microbiota depletion. Notably, in antibiotic-treated, *C. rodentium*–infected mice, fecal pathogen load progressively increased from days 5 to 8 post-infection. However, oral administration of *L. johnsonii* significantly reduced CFUs by day 8 (Fig 6B).

**Fig 6.**
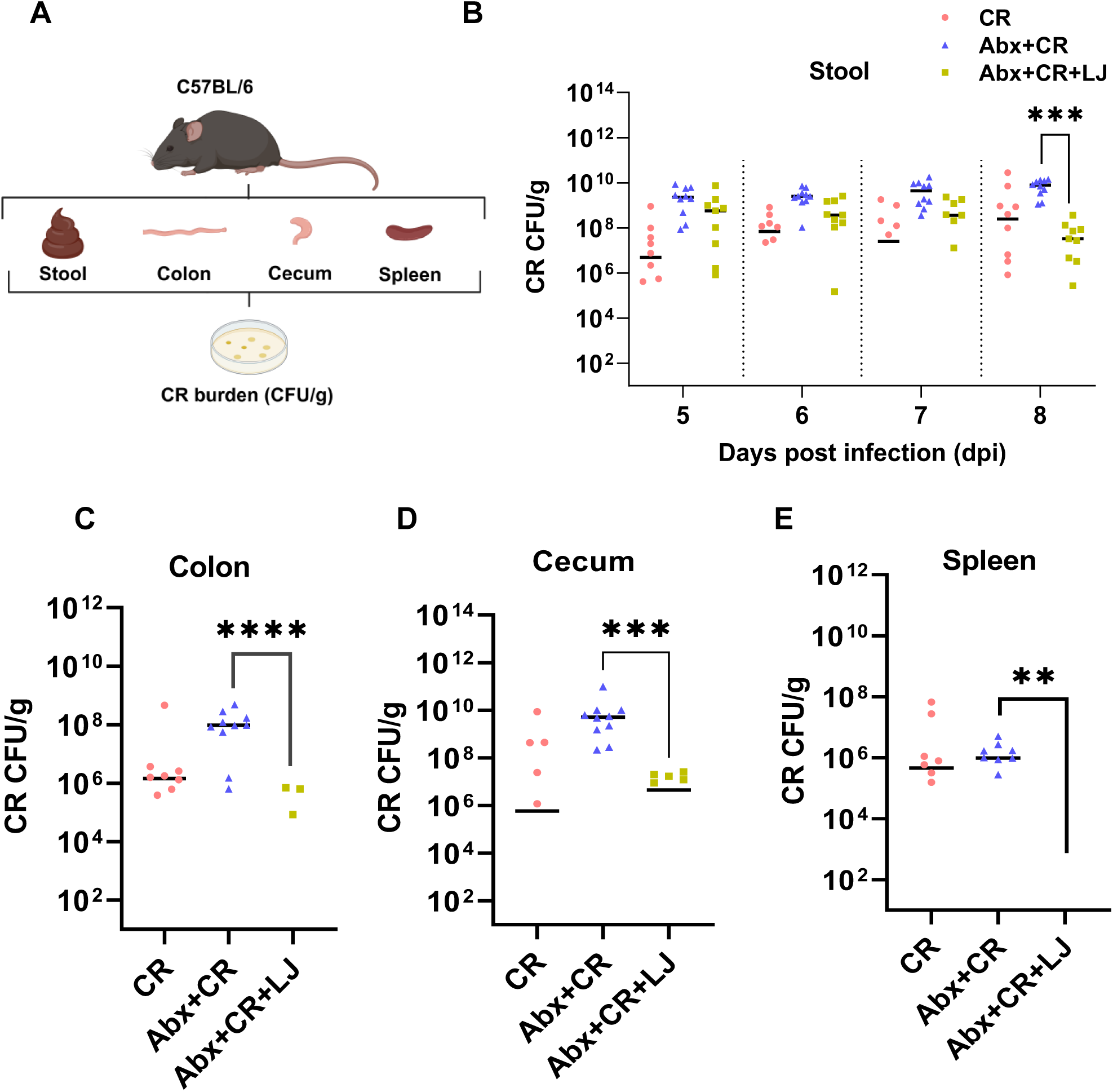
*L. johnsonii* reduces *C. rodentium* burden in stool and tissues. (A) Schematic of the experimental workflow for quantifying *C. rodentium* burden in stool, colon, cecum, and spleen of C57BL/6 mice. (B) Fecal *C. rodentium* loads (CFU/g feces) measured on Days 5-8 post-infection, showing significant reductions in the Abx+CR+LJ group by Day 8 compared to Abx+CR mice. (C-E) Bacterial loads (CFU/g tissue) in (C) colon, (D) cecum, and (E) spleen at Day 8 post-infection. Data are presented as mean ± SEM (n = 9-10 mice/group). Panel B statistics: two-way ANOVA with Dunnett’s multiple comparisons test; \*\**p* < 0.001. Panels C-E statistics: Kruskal-Wallis multiple comparisons test; \**p* < 0.01, \*\**p* < 0.001, \*\*\**p* < 0.0001.

Tissue burden analysis further showed that bacterial titers were substantially higher in the colon (Fig 6C), cecum (Fig 6D), and spleen (Fig 6E) of *L. johnsonii* untreated animals. In contrast, *L. johnsonii* administration led to a highly significant reduction in *C. rodentium* burden across all tissues. These findings demonstrate that *L. johnsonii* not only reduces luminal colonization but also limits mucosal and systemic dissemination, underscoring its potential as a protective probiotic against antibiotic-induced gut vulnerability.

### Nutrient-dependent suppression of *EPEC* and *C. rodentium* by *L. johnsonii*

To investigate the competitive and antimicrobial capacity of *L. johnsonii*, a nutrient competition assay was conducted in McCoy’s 5A medium (nutrient-rich) and phosphate-buffered saline (PBS, nutrient-poor), alongside an agar overlay assay to assess direct inhibitory effects against *C. rodentium* (Fig 7A). In McCoy’s medium, co-culture of *L. johnsonii* with EPEC led to a marked reduction in EPEC CFU (Fig 7B), accompanied by enhanced growth of *L. johnsonii* (Fig 7C), indicating that nutrient competition favors *L. johnsonii*. However, in PBS, no significant changes were observed in either EPEC or *L. johnsonii* growth (Figs 7D–E), highlighting that inhibition is nutrient-dependent. A similar pattern was observed when *L. johnsonii* was co-cultured with *C. rodentium*. In McCoy’s medium, *C. rodentium* growth was significantly suppressed (Fig 7F), while *L. johnsonii* showed enhanced proliferation (Fig 7G). In PBS, no significant differences were noted in CFU counts for either organism (Figs 7H–I), supporting the notion that nutrient availability plays a critical role in the inhibitory interaction. To determine whether *L. johnsonii* also exerts contact-independent killing, an agar overlay assay was performed. Zones of inhibition revealed robust antimicrobial activity of *L. johnsonii* against *C. rodentium*, with greater suppression than gentamicin (Fig 7J), and the inhibition zone was significantly larger in the *L. johnsonii*–treated group (Fig 7K). Collectively, these results suggest that *L. johnsonii* suppresses enteric pathogens through both nutrient-based competition and diffusible antimicrobial factors.

**Fig 7.**
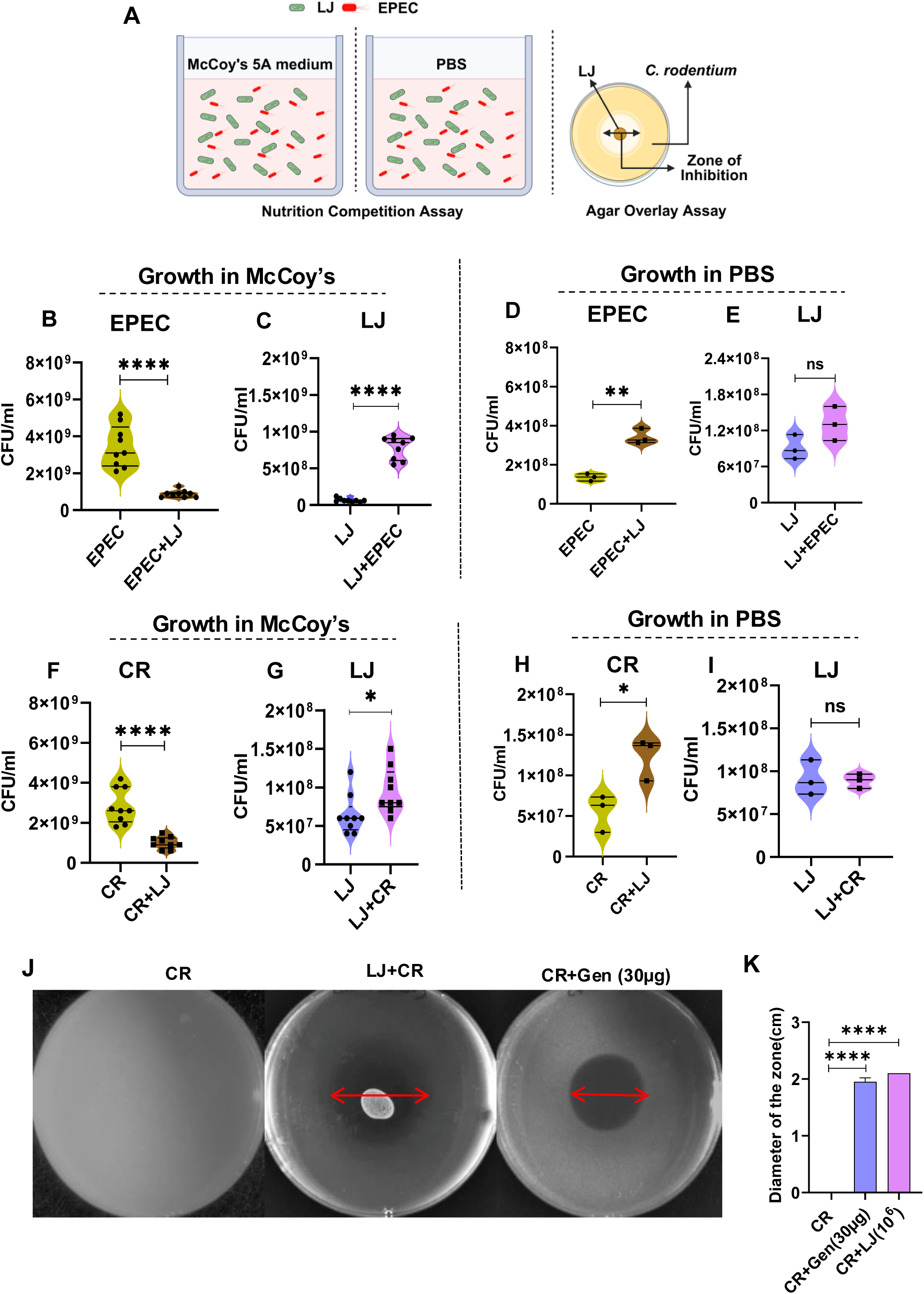
*L. johnsonii* inhibits *EPEC* and *C. rodentium* through nutrient competition and direct antimicrobial activity. (A) Schematic of nutrient competition assay performed in nutrient-rich McCoy’s 5A medium and nutrient-poor phosphate-buffered saline (PBS), and agar overlay assay to assess direct antagonism between *L. johnsonii* (LJ) and EPEC or CR. (B-C) CFU/mL of EPEC (B) and LJ (C) after 6 h co-incubation in McCoy’s 5A medium. (D-E) CFU/mL of EPEC (D) and LJ (E) after 6 h co-incubation in PBS. (F-G) CFU/mL of CR (F) and LJ (G) after 6 h co-incubation in McCoy’s 5A medium. (H-I) CFU/mL of CR (H) and LJ (I) after 6 h co-incubation in PBS. (J) Representative agar overlay plates showing inhibition of CR by gentamicin (30 µg/mL) versus LJ. (K) Quantification of inhibition zone diameters (cm) from agar overlay assay. Data are mean ± SEM from triplicate experiments independently repeated three times (n = 3). Statistical analysis: Student’s *t*-test; \**p* < 0.05; \*\*\*\**p* < 0.0001; ns-not significant.

### Direct antimicrobial effects of *L. johnsonii* against enteropathogenic *E. coli*

To identify bioactive components underlying the antimicrobial activity of *L. johnsonii*, bacterial lysate and culture supernatant were prepared and tested against EPEC as shown in the experimental workflow (Fig 8A). Both treatments significantly reduced EPEC viability, although the lysate exhibited stronger inhibition at lower concentrations (≥12.5 µg), while the cell-free supernatant (CFS) required 50 µg to elicit a comparable effect (Figs 8B–C). To further resolve active components, the CFS was fractionated by FPLC, yielding six major fractions (S1–S6) with molecular weights below 75 kDa (Fig 8D). Antimicrobial activity screening revealed that fractions S1, S2, S5, and S6 significantly inhibited EPEC growth after 1 hour of incubation (Fig 8E). Notably, only fraction S6 retained its inhibitory activity even after 6 hours, indicating the presence of a stable and potent bioactive molecule (Fig 8F). These results demonstrate that *L. johnsonii* exerts direct antimicrobial effects via secreted factors, some of which can be isolated and retained through FPLC, offering potential for future therapeutic development.

**Fig 8.**
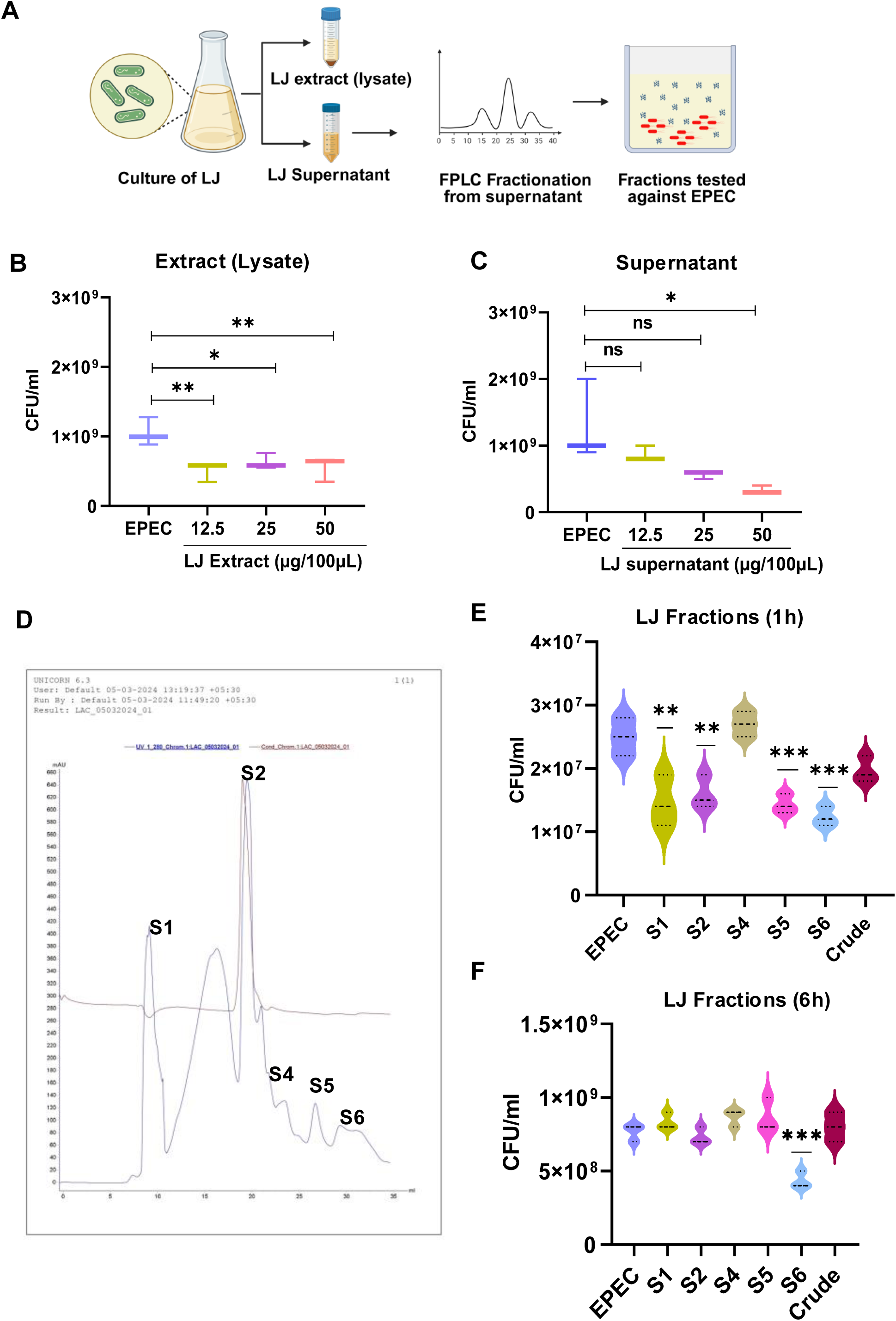
Direct antimicrobial activity of *L. johnsonii* and identification of bioactive FPLC fractions. (A) Experimental workflow for preparing *L. johnsonii* lysate and cell-free supernatant (CFS), followed by fast protein liquid chromatography (FPLC)-based fractionation and antimicrobial testing against EPEC. (B-C) Growth inhibition of EPEC after 6 h treatment with increasing concentrations (12.5, 25, or 50 µg protein/100 µL) of *L. johnsonii* lysate (B) or CFS (C). (D) Representative FPLC chromatogram of *L. johnsonii* CFS showing six eluted fractions (S1-S6), all < 75 kDa in molecular weight, collected using a Superdex 75 size-exclusion column. (E) Antimicrobial activity of individual FPLC fractions (30 µg protein/mL) against EPEC after 1 h incubation. (F) Sustained growth inhibition of EPEC by fraction S6 after 6 h incubation. Data represent mean ± SEM from three independent experiments (n = 3). Statistical analysis: one-way ANOVA with Dunnett’s multiple comparisons test; (\**p* < 0.05; \*\**p* < 0.01; ****p < 0.0001).

### Identification of metabolite classes enriched in *L. johnsonii* antimicrobial fraction S6

To delineate the bioactive compounds responsible for the antimicrobial activity of *L. johnsonii*, we performed metabolomic profiling of the FPLC-purified S6 fraction, which showed the most sustained inhibition of EPEC. Protein staining using SDS-PAGE and Tricine SDS-PAGE revealed no detectable bands in this fraction, ruling out proteinaceous effectors and prompting a small-molecule-based investigation. Mass-spectrometry-based profiling, followed by metabolite set enrichment, revealed that fraction S6 is enriched in fatty acyls, hydroxy acid derivatives, and carboxylic acids that have been associated with membrane disruption and antimicrobial activity (Fig 9A).

**Fig 9.**
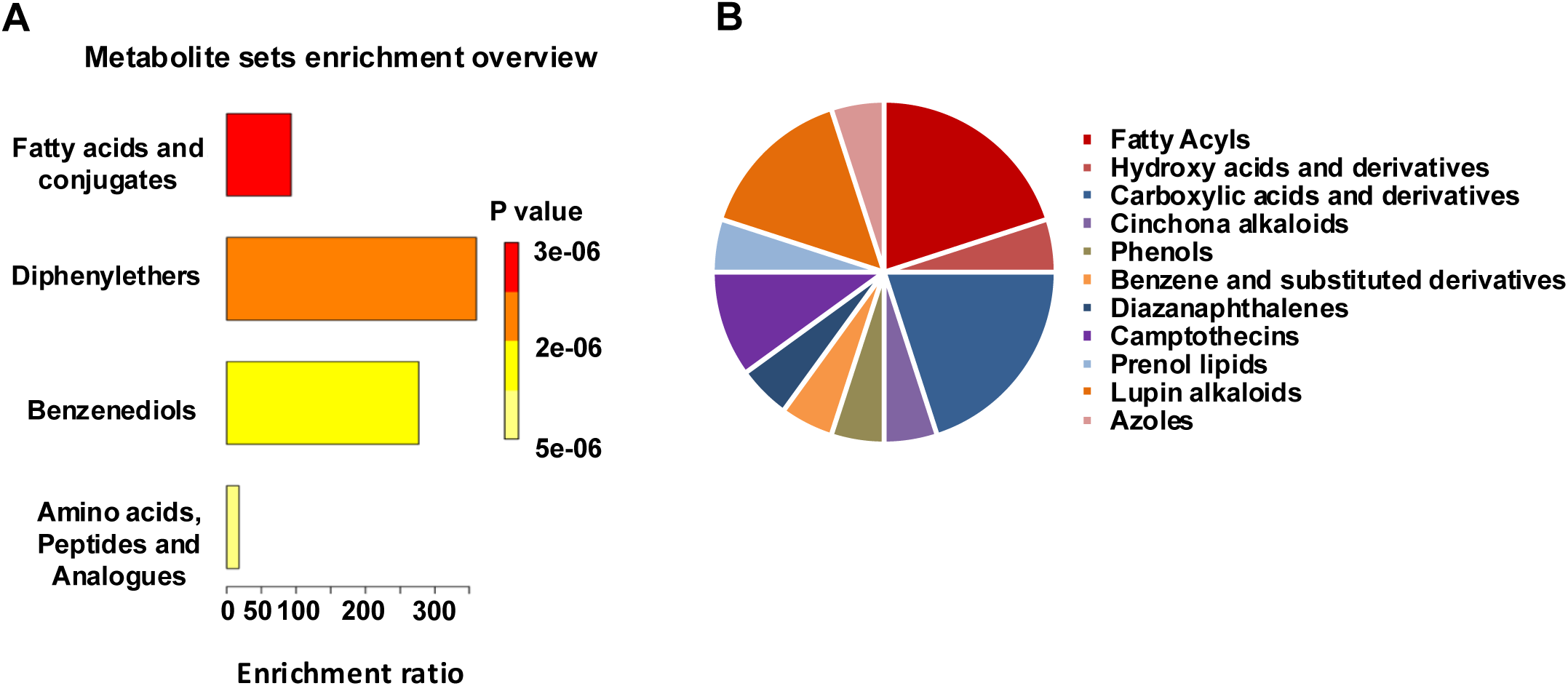
Metabolite profiling of bioactive antimicrobial fraction S6 from *L. johnsonii*. (A) Metabolite set enrichment analysis of the FPLC-purified S6 fraction from *L. johnsonii* cell-free supernatant, showing significant enrichment of antimicrobial-associated chemical classes, including fatty acyls, hydroxy acid derivatives, carboxylic acids, and diphenylethers. Bars represent enrichment ratio with corresponding p-values. (B) Chemical class distribution of 20 metabolites identified by mass spectrometry in positive and negative ion modes, with predominance of fatty acids, hydroxy acid derivatives, alkaloids, and azoles. Of these, 12 putative metabolites have reported antimicrobial activity, and 4 are specifically active against *Escherichia coli* (see S4 Table for details). SDS-PAGE and Tricine SDS-PAGE confirmed no detectable proteins in S6, indicating that antimicrobial activity is mediated by small-molecule metabolites rather than proteinaceous factors.

A complementary chemical class distribution analysis demonstrated a predominance of fatty-acid-related molecules, with notable representation from phenolic compounds, azoles, and alkaloid subclasses including camptothecins and lupin alkaloids (Fig 9B). Of the 20 unique metabolites annotated from positive and negative ionization modes, 12 putative compounds have reported antimicrobial activity, including 4 specifically active against *Escherichia coli* (S4 Table). Among the annotated metabolites, quinine hydrochloride, aloperine, γ-glutamylglutamine, and 1-benzylimidazole have been specifically reported to inhibit *E. coli*. Additionally, high-confidence lipid-based molecules such as 1-linoleoylglycerol, DL-β-hydroxybutyric acid, and decanoic acid are known to perturb bacterial membranes, suggesting multiple complementary antimicrobial mechanisms in the active fraction. These results suggest that the antimicrobial efficacy of *L. johnsonii* is likely mediated by a chemically diverse repertoire of small molecules, with both bacteriostatic and bactericidal potential. The chemical nature and target specificity of these candidate metabolites provide strong rationale for their future isolation and mechanistic validation.

## Discussion

This study establishes *L. johnsonii* as a highly promising probiotic strain with a distinct combination of physiological resilience, antimicrobial potency, and therapeutic efficacy. Unlike widely studied probiotics such as LGG and *L. plantarum*, *L. johnsonii* demonstrated superior performance in both in vitro and in vivo models of enteropathogenic infection, particularly against EPEC and *C. rodentium*. LGG and *L. plantarum* were included as reference strains due to their established clinical use as probiotics for managing pediatric diarrheal diseases, including infections caused by enteric pathogens such as EPEC.

Its enhanced growth in cysteine-supplemented media underscores cysteine’s role as a key metabolic stimulant, likely due to its role as a reducing agent and a precursor for glutathione synthesis, thereby promoting cellular redox balance and metabolic activity [23, 24]. This metabolic flexibility is essential for probiotic persistence under the nutrient-variable conditions of the gastrointestinal tract (GIT), consistent with previous reports on the growth-promoting effects of cysteine in lactobacilli [25, 26]. The robust acid and bile tolerance of *L. johnsonii* further highlights its potential for gut colonization and survival. The GIT presents a challenging environment for microbial survival, characterized by low pH in the stomach and high bile salt concentrations in the small intestine. The ability of *L. johnsonii* to maintain viability at pH levels as low as 1.5 for up to 1 hour and at pH 2.5 for up to 3 hours demonstrates its tolerance to acidic conditions [27]. Similarly, its tolerance to 0.3 % bile salts for at least 3 hours suggests that *L. johnsonii* can withstand the harsh conditions of the upper GIT, a critical requirement for probiotics intended for oral administration [28, 29]. Furthermore, the resistance of *L. johnsonii* to digestive enzymes such as pepsin and trypsin underscores its ability to survive gastrointestinal transit, a key characteristic for probiotics intended for human health applications [30].

The antimicrobial capacity of *L. johnsonii* against EPEC is a key highlight of this study. In agar overlay assays, *L. johnsonii* produced larger zones of inhibition than gentamicin, indicating the secretion of potent antimicrobial compounds. While proteinaceous bacteriocins and organic acids are common antimicrobial effectors among lactobacilli [11, 31], our fractionation results indicate that the predominant bioactive components in *L. johnsonii* are chemically diverse, non-protein small molecules. This was supported by the retention of activity in protein-free FPLC fractions and the MS detection of low-molecular-weight metabolites (<75 kDa). Untargeted metabolomics identified compounds with documented antimicrobial properties, including quinine hydrochloride, aloperine, γ-glutamylglutamine, and 1-benzylimidazole, which have been specifically reported to inhibit *E. coli*. Lipid-based metabolites such as 1-linoleoylglycerol, decanoic acid, and DL-β-hydroxybutyric acid, known to disrupt bacterial membranes, were also detected, suggesting multiple complementary killing mechanisms. Importantly, *L. johnsonii* disrupted pre-formed EPEC biofilms that typically resist both host immune responses and antibiotic treatment [32, 33]. In our study, *L. johnsonii* reduced EPEC biofilm biomass by ∼60 %, markedly higher inhibition than observed with reference strains (LGG, *L. plantarum*), highlighting its potent biofilm-disrupting capacity [34–36]. The partial retention of inhibitory activity with heat-killed cells suggests that at least some antimicrobial effectors are heat-stable, consistent with our identification of small-molecule metabolites as major active agents. This reduces the likelihood that structural cell wall components are the dominant inhibitory factor [16, 17, 37].

Nutrient competition assays revealed both competitive and direct antimicrobial effects. In nutrient-rich environments, *L. johnsonii* significantly suppressed the growth of EPEC and *C. rodentium*, indicating its ability to outcompete these pathogens for essential nutrients. Conversely, this inhibitory effect was attenuated under nutrient-limited conditions, highlighting the importance of nutrient availability in mediating its antagonistic activity. These findings are consistent with the principle of nutritional immunity, wherein commensal microbes and pathogens compete for critical micronutrients such as iron, zinc, and amino acids to establish dominance within the host environment [38]. Notably, the bacterial lysate exhibited significant inhibitory activity even at 12.5 µg, while the CFS required a higher concentration of 50 µg to achieve similar suppression, indicating that intracellular components may contribute more potently to EPEC inhibition than secreted factors alone.

Subsequent fractionation of the *L. johnsonii* cell-free supernatant identified multiple fractions with antimicrobial activity, with fraction S6 showing sustained inhibition of EPEC growth. SDS-PAGE and Tricine SDS-PAGE analysis of S6 failed to reveal detectable protein bands, prompting metabolomic profiling of this fraction. The resulting analysis revealed a diverse repertoire of small molecules, including fatty acids, hydroxy acid derivatives, and alkaloid-like compounds known to exert antimicrobial and membrane-disruptive effects. Several detected metabolites, including quinine hydrochloride dihydrate and rubitecan, have not previously been reported in probiotic-derived fractions, highlighting the unexpected chemical diversity of compounds secreted by *L. johnsonii*. While the precise contribution of individual metabolites remains to be validated, their chemical properties are consistent with known membrane-targeting antimicrobials. This aligns with Fig 9, which identifies fatty acyls and alkaloids, both implicated in disrupting bacterial membranes or interfering with quorum sensing as dominant constituents of the active fraction. These findings provide a strong foundation for future bio-guided isolation and structural characterization of *L. johnsonii*-derived metabolites, which may hold translational promise as next-generation antimicrobials against drug-resistant enteric pathogens.

Beyond biofilm inhibition, *L. johnsonii* also demonstrated a distinct advantage in disrupting pathogen adhesion to host epithelium. LGG, for instance, employs mucus-binding pili (SpaCBA) to adhere to the intestinal mucosa and inhibit initial pathogen attachment [39, 40] but this mechanism is less effective once pathogens are already bound to the epithelial surface [40]. In displacement assays, *L. johnsonii* removed over 50 % of pre-adhered EPEC, showing greater efficacy than reference strains, particularly *L. plantarum*, which adhered poorly to HCT-116 cells. This activity may be mediated by surface-layer (S-layer) proteins and fibronectin-binding domains that disrupt pathogen-host interactions, features not reported in LGG and minimally observed in *L. plantarum* [41]. Although *L. plantarum* can adhere to mucosal surfaces in other models, it showed poor adhesion to HCT-116 cells and was markedly less effective at displacing EPEC [42]. Notably, while *L. johnsonii* did not significantly prevent initial pathogen binding, it consistently disrupted established EPEC colonization, suggesting a mechanism based on competitive exclusion and interference with biofilm integrity [43].

Our findings on the antibiotic resistance profile of *L. johnsonii*, particularly its resistance to kanamycin and gentamicin, are consistent with intrinsic resistance traits commonly observed in *Lactobacillus* species. Importantly, these resistance genes appear to be chromosomally encoded rather than plasmid-borne, thereby reducing the risk of horizontal gene transfer to pathogenic bacteria [44]. This is supported by earlier work from Danielsen and Wind (2003), who showed that *Lactobacillus* spp. rarely transfer resistance genes under physiological conditions [20]. This intrinsic resistance, coupled with its susceptibility to non-prophylactic antibiotics like ampicillin, positions *L. johnsonii* as a safe probiotic for clinical use, even in antibiotic-exposed populations [45].

The murine model experiments provide compelling evidence for *L. johnsonii*’s protective role against *C. rodentium*-induced colonic pathology. *C. rodentium* is a natural murine pathogen that shares key virulence factors with EPEC and enterohemorrhagic *E. coli*, making it a well-established model for studying attaching and effacing (A/E) lesion-forming pathogens [46]. In our study, antibiotic treatment effectively eradicated *L. johnsonii* and significantly reduced total gut bacterial load, underscoring the broad-spectrum impact of antibiotics on microbiota composition and the increased susceptibility to enteric pathogens reported previously [47, 48]. The loss of colonization by *L. johnsonii* and depletion of microbial diversity likely created an ecological niche favorable for *C. rodentium* overgrowth [49, 50]. Therapeutic administration of *L. johnsonii* following antibiotic treatment markedly mitigated colonic inflammation, mucosal edema, and pathological damage, demonstrating its capacity to restore gut homeostasis. These benefits were accompanied by restoration of colon length and reduced inflammatory scores, consistent with prior studies showing that probiotics can attenuate loss of barrier integrity [51, 52]. Our findings align with earlier reports that *L. johnsonii* promotes epithelial cell proliferation and reduces inflammatory signaling in intestinal epithelial cells [53–55]. Additionally, the observed reduction in neutrophil infiltration in *L. johnsonii* treated mice supports the hypothesis that this strain can downregulate pro-inflammatory cytokine production, thereby limiting tissue damage and facilitating recovery [56]. Collectively, these results demonstrate that *L. johnsonii* exerts multifaceted protective effects in an A/E pathogen infection model, consistent with its in vitro antimicrobial and anti-biofilm activities.

Our study has limitations. The active S6 fraction was defined phenotypically; specific molecules still need confirmation with authentic standards (MSI Level 1) and purification for direct activity testing. Epithelial findings were generated in a single non-mucus cell line and at a single MOI selected on the basis of cytotoxicity experiments, and in vivo efficacy was demonstrated in an antibiotic-sensitized mouse model. Broader cellular models including mucus-producing or pri mary epithelia, alternative dosing regimens, and targeted chemical validation will deepen mechanistic understanding and broaden applicability.

## Conclusion

In conclusion, *L. johnsonii* combines robust gastrointestinal resilience, potent small-molecule-mediated antimicrobial activity, biofilm disruption, pathogen displacement, and mitigation of infection-induced colonic damage. Its efficacy in post-infection and antibiotic-compromised settings offers a compelling non-antibiotic strategy to address the urgent challenge of antimicrobial resistance. Future studies should prioritize mechanistic dissection of its active components, explore interactions with the host microbiota, and evaluate clinical potential, particularly in pediatric populations disproportionately affected by antibiotic-resistant diarrheal disease.

## Materials and Methods

### Bacterial strains, media and growth conditions

*L. johnsonii* was isolated from homemade Indian curd collected in Hyderabad, India. The isolate was cultured on de Man, Rogosa, and Sharpe (MRS) broth (Millipore Sigma, Cat. No. 69966) supplemented with 0.05% (w/v) cysteine (Sigma-Aldrich, Cat. No. 168149) for 12 hours at 37°C and its identity was confirmed by 16S rRNA gene sequencing. *L. plantarum* (NCDC-NDRI) *Enteropathogenic Escherichia coli* (EPEC) strain was provided by Dr. Tannaz J. Biridi (Foundation for Medical Research, Mumbai, India), and *C. rodentium* (ATCC 51459) was obtained from Dr. Soumen Basak (National Institute of Immunology, India). EPEC and *C. rodentium* were grown in Luria-Bertani (LB) broth (Himedia, Cat. No. M1245) at 37°C at 200 rpm.

### Growth curve analysis and colony-forming unit (CFU) determination

*L. johnsonii* and *L. plantarum* were cultured in 10 mL of MRS broth or MRS supplemented with 0.05% (w/v) cysteine (MRS+), while EPEC was grown in 10 mL of LB broth. All cultures were incubated at 37°C with shaking at 200 rpm. Optical density at 600 nm (OD₆₀₀) was measured at 0, 2, 4, 8, 24, 48, and 72 hours for *Lactobacillus* strains, and up to 24 hours for EPEC, using a BioSpectrometer® Basic (Molecular Devices). For CFU determination, overnight cultures of *L. johnsonii*, EPEC, and *C. rodentium* (adjusted to OD₆₀₀ = 1) were serially diluted tenfold (10⁻¹ to 10⁻⁷). From each dilution, 10 µL was plated in triplicate on LB agar (for EPEC and *C. rodentium*) or MRS+ agar (for *L. johnsonii*) and incubated overnight at 37°C. Viable counts were calculated as:

CFU/mL = (Number of colonies × dilution factor) / volume plated.

### Preparation of artificial gastrointestinal fluids and assessment of bacterial tolerance

*L. johnsonii* was incubated in MRS^+^broth adjusted to pH 1.5-2.5 (using 1 M HCl) or supplemented with 0.3% (w/v) bile (Himedia-CR010) for 0-3 hours. Viability was assessed by plating on MRS^+^agar and calculating CFU/mL. Gastric fluid was prepared by dissolving pepsin (3.0 g/L; Lonza) in sterile phosphate-buffered saline (PBS), followed by pH adjustment to 2.5 using 0.1 N hydrochloric acid. The solution was sterilized by filtration through a 0.22-μm membrane filter (Sartorius). Intestinal fluid was prepared by dissolving trypsin (1.0 g/L; SRL) and bile salts (1.8% w/v; HiMedia) in sterile PBS, followed by pH adjustment to 8.0 with 0.1 M sodium hydroxide and subsequent filtration through a 0.22-μm membrane. Bacterial cultures (10⁶ CFU/mL) were harvested by centrifugation, washed three times with sterile PBS, and resuspended to a final concentration of 0.1 mL. This suspension was added to 1 mL of either artificial gastric or intestinal fluid and incubated at 37°C for 3 hours under anaerobic conditions. Viable bacterial counts were determined post-incubation by serial dilution and plating, and results were expressed as CFU/mL.

### Agar overlay assay

*L. johnsonii* was spot-inoculated onto MRS^+^agar plates using 10 μl (approximately 10⁵ CFU/spot) of MRS^+^broth culture and incubated at 37°C for 48 hours. Following incubation, the MRS^+^agar plates with *L. johnsonii* growth were overlaid with LB agar containing 10⁷ CFU of EPEC per plate. After solidification of the overlaid agar, the plates were incubated at 37°C for 24 hours. The resulting zone of inhibition (ZOI) was measured and compared with the ZOI of gentamicin.

### Biofilm formation assay

To evaluate inhibition of EPEC biofilm formation in the presence of *L. johnsonii*. EPEC and *L. johnsonii* were inoculated individually, as well as together, into the wells of a 96-well plate containing Brain Heart Infusion (BHI) broth. After 48 hours of incubation, the liquid was removed, and the wells were washed three times with distilled water. The biofilm was then stained with 0.1% crystal violet for 30 minutes, followed by three additional washes with distilled water. The plate was air-dried, and 100 µl of 100% ethanol was added to each well to solubilize the bound crystal violet. The absorbance of the resulting solution was measured at 570 nm using a spectrophotometer (Molecular devices).

### Multiplicity of Infection (MOI) determination on HCT-116 Cells

HCT-116 cells (NCCS) were maintained in McCoy’s 5A medium (Gibco, Cat. No.16600082) combined with 10% FBS (Gibco-10270-106). HCT-116 is a non-mucus-producing human intestinal epithelial cell line, making it well-suited for direct bacteria-epithelial interaction studies without interference from mucus layers. Cell monolayers were infected with *L. johnsonii* or EPEC at MOIs of 1:15 to 1:100 (cell: bacteria). After 6 hours, cytotoxicity was quantified from supernatants using an LDH assay kit (Thermo Fisher, Cat. No. 88953) according to the manufacturer’s instructions

### Antibiotic susceptibility assays

The antibiotic susceptibility of *L. johnsonii* and *L. plantarum* to ampicillin, kanamycin, gentamicin, and vancomycin was assessed by testing these antibiotics at concentrations recommended by the European Committee on Antimicrobial Susceptibility Testing (EUCAST). Bacterial cultures (10⁶ CFU/mL) were subjected to varying concentrations of antibiotics in 96-well plates as noted in S2 Table. Uninoculated MRS^+^broth and untreated bacterial cultures served as negative and positive controls, respectively. After a 24-hour incubation at 37°C, 10 µL aliquots from each well were spot-plated onto MRS^+^ agar and incubated overnight at 37°C to evaluate bacterial viability.

### Adherence and protection Assays

To assess the adherence of probiotics (*L. johnsonii* and *L. plantarum*) and *EPEC* to HCT-116 cells, monolayers were seeded in 24-well plates (2.5 × 10⁴ cells/well) and grown to 90% confluence. Cells were incubated with bacteria at an MOI of 1:25 for 3 hours. After washing with PBS to remove non-adherent bacteria, cells were lysed with 1% Triton X-100, and serial dilutions were plated on MRS^+^agar for *L. johnsonii* and *L. plantarum*, and LB agar for *EPEC*. Plates were incubated at 37°C overnight, and CFUs were enumerated. For protection assays, HCT-116 cells grown to 90% confluence were exposed to probiotics and EPEC (MOI 1:25) under three conditions: (i) Exclusion Assay: Cells were pre-incubated with *L. johnsonii* or *L. plantarum* for 3 hours before infecting with EPEC for another 3 hours. (ii) Displacement Assay: Cells were infected with EPEC for 3 hours, after which *L. johnsonii* or *L. plantarum* was added for another 3 hours. (iii) Competition Assay: *L. johnsonii* or *L. plantarum* and EPEC were co-incubated for 6 hours. After incubation, wells were washed three times, and adherent bacteria were quantified by lysing cells with 1% Triton X-100. Serial dilutions were plated on MRS^+^agar for *Lactobacillus* and LB agar for EPEC, followed by incubation at 37°C for CFU enumeration.

### Mice, antibiotic treatment, *C. rodentium* infection and probiotic treatment

Six-week-old female C57BL/6 mice, procured from the National Institute of Nutrition, were housed in the animal facility at the University of Hyderabad (UoH) in accordance with Institutional Animal Ethics Committee (IAEC) guidelines (UH/IAEC/VM/2021-1/28) and the Committee for the Purpose of Control and Supervision of Experiments on Animals (CPCSEA), India. Mice were maintained under controlled environmental conditions (22 ± 2 °C temperature, 50–60% relative humidity, 12 h light/12 h dark cycle) with standard cage enrichment and ad libitum access to food and water. Antibiotics were selected to broadly deplete Gram-positive, Gram-negative, and anaerobic bacteria, mimicking clinical dysbiosis. Mice received a single oral gavage (200 μL) of an antibiotic cocktail containing neomycin, gentamicin, metronidazole, and streptomycin (each at 2 mg/mL) and vancomycin (1 mg/mL) (HiMedia). This was followed by a seven-day regimen in which drinking water was supplemented with neomycin, gentamicin, metronidazole, and streptomycin (each at 1 mg/mL), vancomycin (0.5 mg/mL), and 10% sucrose [57]. On day 8, mice were orally gavaged with 100 µl of PBS containing *C. rodentium* at a concentration of 2.5 × 10⁸ CFU per mouse. From days 9 to 13, the probiotic-treated group received *L. johnsonii* (10⁹) CFU/100ul/ mouse). Control groups included untreated mice, *C. rodentium*-infected mice without antibiotics, and antibiotic-treated mice infected with *C. rodentium*. Mice were randomly assigned to experimental groups, and investigators were blinded to group allocation during outcome assessment. The sample size (n = 9-10 per group) was determined based on effect sizes observed in previous *C. rodentium* infection studies using C57BL/6 mice, ensuring adequate statistical power while adhering to the principle of reduction in animal use. The primary endpoint was the reduction in *C. rodentium* burden in feces and colonic tissue at 8 days post-infection, with secondary endpoints including colon length, histopathological scores, and inflammatory cell infiltration. Baseline body weights were recorded before infection, and mice were weighed daily to monitor changes. Weight variations were normalized and plotted as percentage changes relative to baseline values. On day 8 post-infection, mice were euthanized and tissues were aseptically collected for further analysis.

### Fecal genomic DNA isolation, Gel electrophoresis and quantitative real-time PCR

Genomic DNA was extracted from the collected fecal samples from all the mice groups on Day 5, 6, 7 and 8 post infection using a DNA isolation kit (GCC-BIOTECH). The DNA concentration was then measured using a NanoDrop Spectrophotometer (Thermo Scientific). Each PCR reaction mixture consisted of 25 µl of PCR Master Mix, 0.2 µl of both forward and reverse primers, and 5 µl of template. PCR was performed in a thermal cycler under the following conditions: initial denaturation at 95°C for 5 minutes, followed by 40 cycles of denaturation at 95°C for 10 seconds, annealing at 60°C for 30 seconds, and extension at 72°C for 40 seconds, with a final extension step at 72°C for 5 minutes. Following amplification, the PCR products were analyzed using electrophoresis on a 1.5% agarose gel and visualized under vilber gel imaging system. Quantitative real-time PCR (qPCR) was performed using a Qiagen qPCR kit under the same cycling conditions as conventional PCR, with primer sequences listed in Table 1. Total bacterial load was determined by the standard curve method, using genomic DNA from a known number of *Escherichia coli* cells as the calibrator for universal 16S rRNA quantification (Fig 4D). For confirmation of probiotic strain identity, universal 16S rRNA gene primers (27F: 5′-AGAGTTTGATCCTGGCTCAG-3′ and 1492R: 5′-GGTTACCTTGTTACGACTT-3′) were used to amplify an approximately 1.5 kb fragment, which was subsequently sequenced by the Sanger method

**Table 1:**
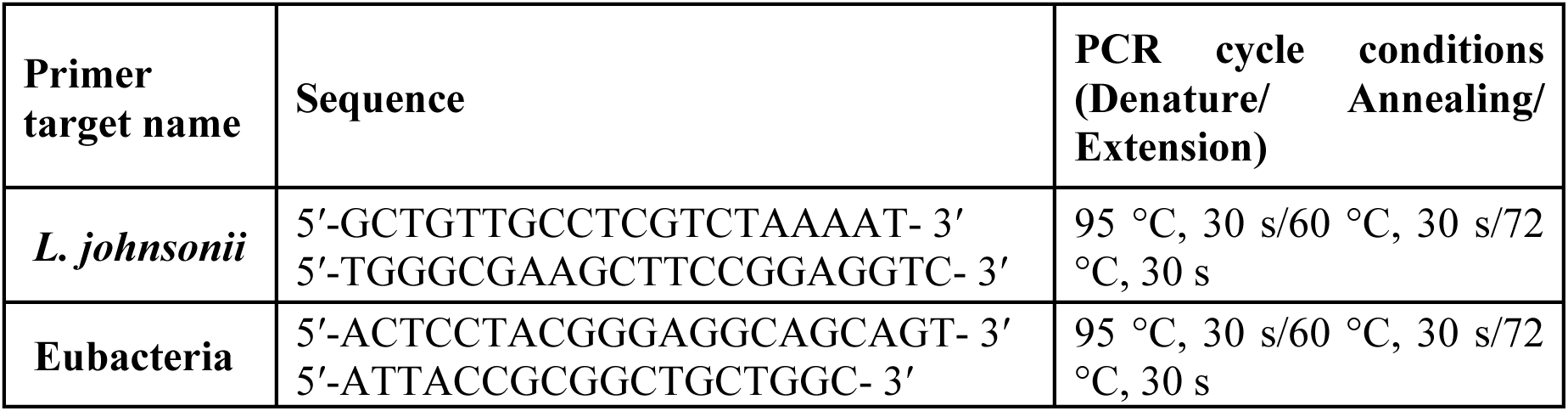
Primer sequences and PCR cycling conditions used for the detection of *L. johnsonii* and total eubacteria.

### CFU counting from mouse tissues and stool samples

The colons, ceca, spleens, and livers of the mice were collected in PBS and homogenized using a tissue homogenizer (Cole-Parmer**®**). Stool samples were collected on day 5, 6, 7 and 8 in 500 µl of PBS and homogenized. Serial dilutions (10^-1^-10^-7^) of the homogenates were plated on LB-Amp agar plates to assess *C. rodentium* colonization. The plates were incubated overnight at 37°C, after which colonies were counted and expressed as CFU per gram of tissue.

### Histopathological scoring

Hematoxylin and eosin (H&E)–stained tissues were scored independently by two blinded observers. Four parameters were assessed: (i) submucosal edema (0 = no change, 1 = mild, 2 = moderate, 3 = severe); (ii) goblet cell depletion (0 = no change, 1 = mild depletion, 2 = severe depletion, 3 = absence of goblet cells); (iii) ulceration, when present, with small ulcers defined as < 25% of the cross-sectional area, medium ulcers as > 25–50%, and large ulcers affecting 75% of the section scored as 4; and (iv) inflammatory cell infiltration per ×400 field (0 = no change, 1 = 50 cells/field). The maximum possible pathology score with this scheme was 20.

### Immunostaining

Formalin-fixed, paraffin-embedded (FFPE) colon tissue sections (5 µm) were deparaffinized at 60 °C for 15 min, cleared in xylene, and rehydrated through a graded ethanol series. Antigen retrieval was performed in 10 mM sodium citrate buffer (pH 6.0) at 95 °C for 45 min. After cooling to room temperature, sections were blocked for 1 h in PBS containing 5% (v/v) goat serum, 1% (w/v) bovine serum albumin (BSA), 0.1% Triton X-100, 0.05% Tween-20, and 0.05% sodium azide to prevent nonspecific binding. Tissues were incubated overnight at 4 °C with the following primary antibodies: rat anti-mouse Ly6G (1:200; Cell Signaling Technology, Cat# 87048, clone RB6-8C5) and rabbit anti-mouse Ki-67 (1:200; Abcam, Cat# ab16667, clone SP6). After three washes in PBS containing 0.05% Tween-20, sections were incubated for 1 h at room temperature in the dark with species-specific fluorophore-conjugated secondary antibodies: Alexa Fluor® 488 goat anti-rat IgG (H+L) (1:400; Invitrogen, Cat# A-11006) and Alexa Fluor® 568 goat anti-rabbit IgG (H+L) (1:400; Abcam, Cat# ab175471). Nuclei were counterstained using ProLong™ Gold Antifade Mountant with DAPI (Invitrogen, Cat# P36931). Sections were mounted with coverslips and visualized using an Olympus IX71 inverted fluorescence microscope equipped with a DP23M high-sensitivity camera, and images were acquired and analyzed using cellSens™ software (Olympus Corporation, Japan).

### Nutrient competition assay

Overnight cultures of *L. johnsonii*, EPEC, and *C. rodentium* were adjusted to 10⁷ CFU/well in a 24-well plate. *L. johnsonii* was centrifuged at 3,000 rpm for 5 minutes, and bacterial pellets were resuspended in infection media (McCoy’s incomplete media + 2.5% FBS). *L. johnsonii* was co-incubated with either EPEC or *C. rodentium* for 6 hours. Post-incubation, 10 µl of serially diluted samples were plated on MRS^+^agar for *L. johnsonii* and LB agar for EPEC and *C. rodentium*. CFU/mL were enumerated after incubation at 37°C.

### Preparation of bacterial cell-free supernatant and lysate

Bacterial cell-free supernatant (CFS) was prepared by culturing *L. johnsonii* in MRS^+^broth for 18-24 hours, followed by centrifugation at 3,500 rpm for 20 minutes. The supernatant was filtered through a 0.22 μm pore-size filter. The pellet was washed with 0.85% NaCl and resuspended in 1X PBS to obtain a final concentration of 1×10⁹ CFU/mL. For bacterial lysate preparation, 10 mL of bacterial suspension was centrifuged and incubated at 37°C for 2 hours in lysozyme solution (10 mg/mL in Tris-EDTA buffer, pH 8.0). The cells were washed, resuspended in 1X PBS, and sonicated using a Sonopulse Probe (25 cycles of 15 seconds each, with rest intervals on ice). Lysates were filtered through a 0.22 µm-pore filter (Millipore). Protein content in CFS and lysate was quantified using a DC^TM^ protein assay (Bio-rad) with bovine serum albumin as the standard.

### Fast protein liquid chromatography (FPLC)

*L. johnsonii* was cultured in 2 L of MRS^+^broth (Millipore Sigma; Cat. No. 69966) at 37°C with agitation at 200 rpm for 48 hours. Following incubation, the culture was centrifuged at 3,500 rpm (Hitachi) and the supernatant was filtered through a 0.22-μm cellulose nitrate membrane (Sartorius). The clarified supernatant was stirred continuously at 8°C overnight. Ammonium sulfate (SRL; Cat. No. 82126) was added to a final concentration of 30% (w/v), and the solution was incubated at 8°C for 18 hours. The precipitate was collected by centrifugation at 12,000 × g for 15 min at 6°C. The resulting surface film was re-centrifuged, resuspended in 50 mM phosphate buffer (pH 7.0), and concentrated by lyophilization (Scanvac) overnight. Approximately 5 mg of protein in 500 µL phosphate buffer was subjected to FPLC using a Superdex 75 size exclusion column (Cytiva). Eluted fractions (S1-S6) were collected, lyophilized, and individually assessed for antimicrobial activity against *Enteropathogenic Escherichia coli* (EPEC).

### Metabolite profiling of antimicrobial FPLC fraction

The bioactive fraction (S6) obtained via FPLC from the *L. johnsonii* cell-free supernatant was subjected to untargeted metabolite profiling by mass spectrometry (MS). A 2 µL aliquot of the S6 fraction was injected into an LCMS-8040 system (Shimadzu) operated at a flow rate of 200 µL/min. Electrospray ionization (ESI) was applied in both positive and negative ion modes to maximize metabolite coverage. Raw data files were converted to Analysis Base File (ABF) format using the ABF converter and processed in MS-DIAL (version X.X). The separation type was set to direct infusion, and MS1 spectra were averaged across scans. Feature alignment was performed using a peak detection threshold of ≥66 % reproducibility across triplicate samples. Metabolite annotation was carried out against public spectral libraries, including MSMS-Pos-MassBank and MSMS-Public_Pos_VS19_1 for positive mode, and MSMS-Neg-MassBank and MSMS-Public_Neg_VS19_1 for negative mode. Spectral matches were further confirmed using the MassBank of North America (MoNA) library. Following metabolite identification, enrichment analysis was performed in MetaboAnalyst (version X.X) to classify compounds into functional chemical groups. The resulting metabolite set enrichment plots and class distribution charts are presented in Fig 9A-B. Annotated metabolites are summarized in S4 Table, with known antimicrobial compounds highlighted.

### Statistical analysis

All biological assay data are presented as mean ± standard deviation (SD) or standard error of the mean (SEM), as indicated in Fig legends. Statistical analyses were performed using GraphPad Prism version 8.4.2 (GraphPad Software, San Diego, USA). Depending on the experimental design, group comparisons were conducted using unpaired or paired two-tailed Student’s t-tests, one-way ANOVA with Tukey’s multiple comparisons test, Kruskal–Wallis test with Dunn’s multiple comparisons, and two-way ANOVA with Dunnett’s multiple comparisons test. For two-group comparisons that did not meet t-test assumptions, the Mann– Whitney U test was used. A p-value < 0.05 was considered statistically significant. Significance thresholds are denoted in Figs as follows: *p* < 0.05 (*)*, < 0.01* (**) ***<*** *0.001* (***), and < 0.0001 (****); “ns” indicates no statistical significance. Unless otherwise specified, the unit of analysis was the individual animal for all in vivo outcomes. For histology and immunostaining, representative images are shown; quantitative analyses were performed with a minimum of three biological replicates per group. For microbiological assays, the unit of analysis was a single tissue sample per animal, and for in vitro assays, the unit of analysis was an individual well or replicate.

## Acknowledgments

The authors gratefully acknowledge Dr. Tannaz J. Biridi (Foundation for Medical Research, Mumbai, India) for providing the *Enteropathogenic Escherichia coli* (EPEC) strain and Dr. Soumen Basak (National Institute of Immunology, India) for generously sharing the *Citrobacter rodentium* (ATCC 51459) strain. We also thank Dr. Prasada Rao H. B. D. from the National Institute of Animal Biotechnology for his valuable insights on FPLC. The authors also acknowledge the technical staff at AIG Hospitals, Hyderabad, for their assistance in preparing colon tissue sections and performing hematoxylin and eosin (H&E) staining as well as immunofluorescence staining. We sincerely appreciate the contributions of laboratory members for their valuable discussions and assistance during the course of this study. The authors reviewed and take full responsibility for the content.

## Supporting Information

### Supplementary Fig Legends

**S1 Fig:**
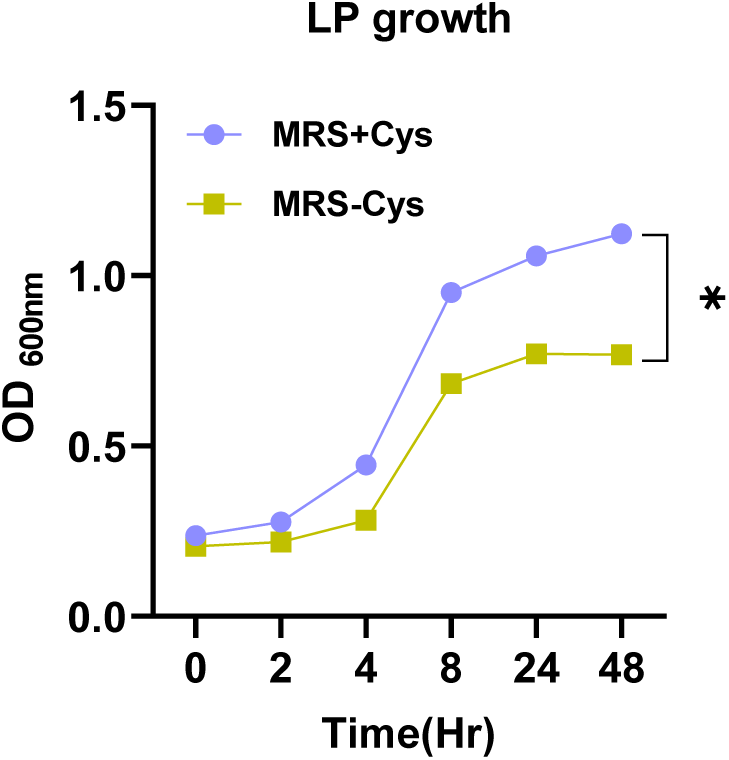
Growth curves of *L. plantarum* in de Man, Rogosa and Sharpe (MRS) broth supplemented with 0.05% (w/v) cysteine (MRS+Cys) versus unsupplemented MRS (MRS-Cys) over 0-72 h incubation at 37 °C. Optical density at 600 nm (OD₆₀₀) was recorded at the indicated time points. Data are mean ± SEM from three independent biological replicates.

**S2 Fig:**
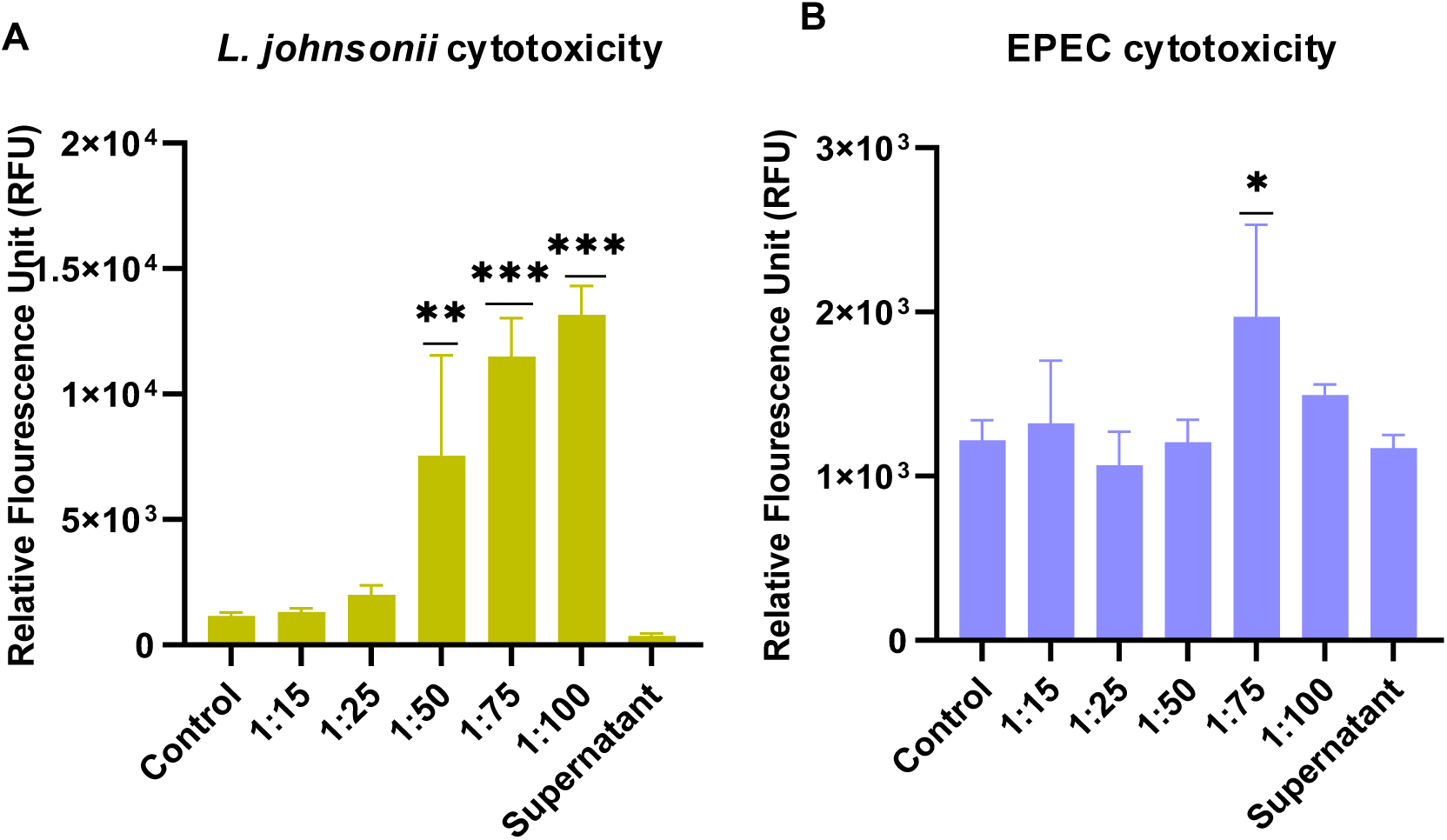
Cytotoxicity induced by *L. johnsonii* and EPEC in HCT-116 cells. (A) Lactate dehydrogenase (LDH) release from HCT-116 intestinal epithelial cells after 6 h exposure to *L. johnsonii* at different multiplicities of infection (MOI). (B) LDH release from HCT-116 cells after 6 h exposure to EPEC at different MOIs. LDH release (%) was calculated relative to maximum release induced by cell lysis (positive control) and untreated cells (negative control). Data represent the mean ± SEM from two independent experiments performed in triplicate. Statistical analysis: one-way ANOVA \**p* < 0.05; \*\**p* < 0.01; \*\*\**p* < 0.001.

**S3 Fig:**
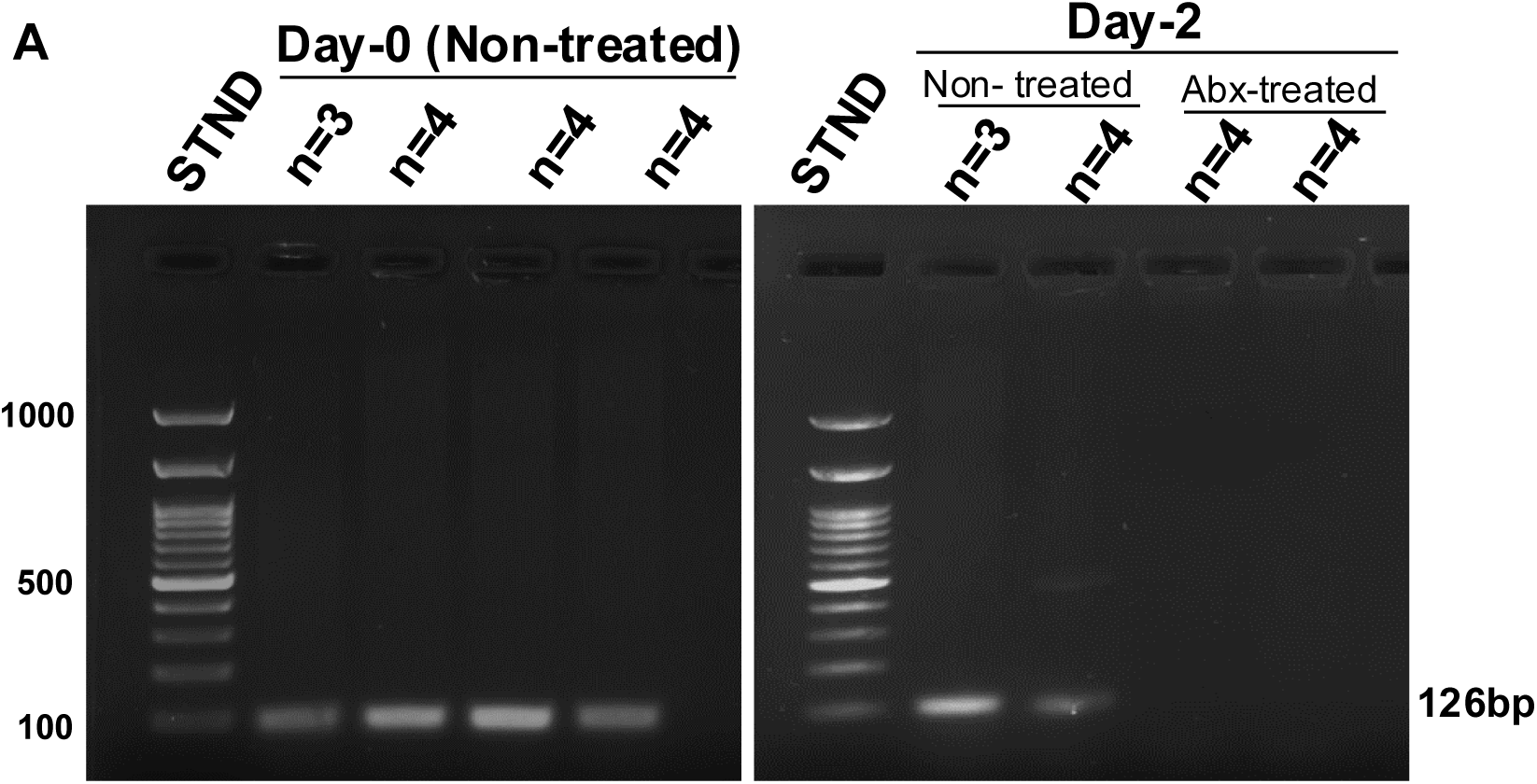
Early depletion of *L. johnsonii* following antibiotic treatment. (A) Agarose gel electrophoresis showing absence of the *L. johnsonii*-specific 100 bp PCR amplicon in pooled stool DNA from antibiotic-treated mice by Day 2. DNA was extracted from stool samples of n = 4 mice per group and pooled before analysis. Lane 1 and Lane 6: 100 bp DNA ladder; Lanes 2-5: pooled stool DNA samples from Day 0 (pre-antibiotic treatment); Lanes 7-10: pooled stool DNA samples from Day 2 post-antibiotic treatment.

**S4 Fig:**
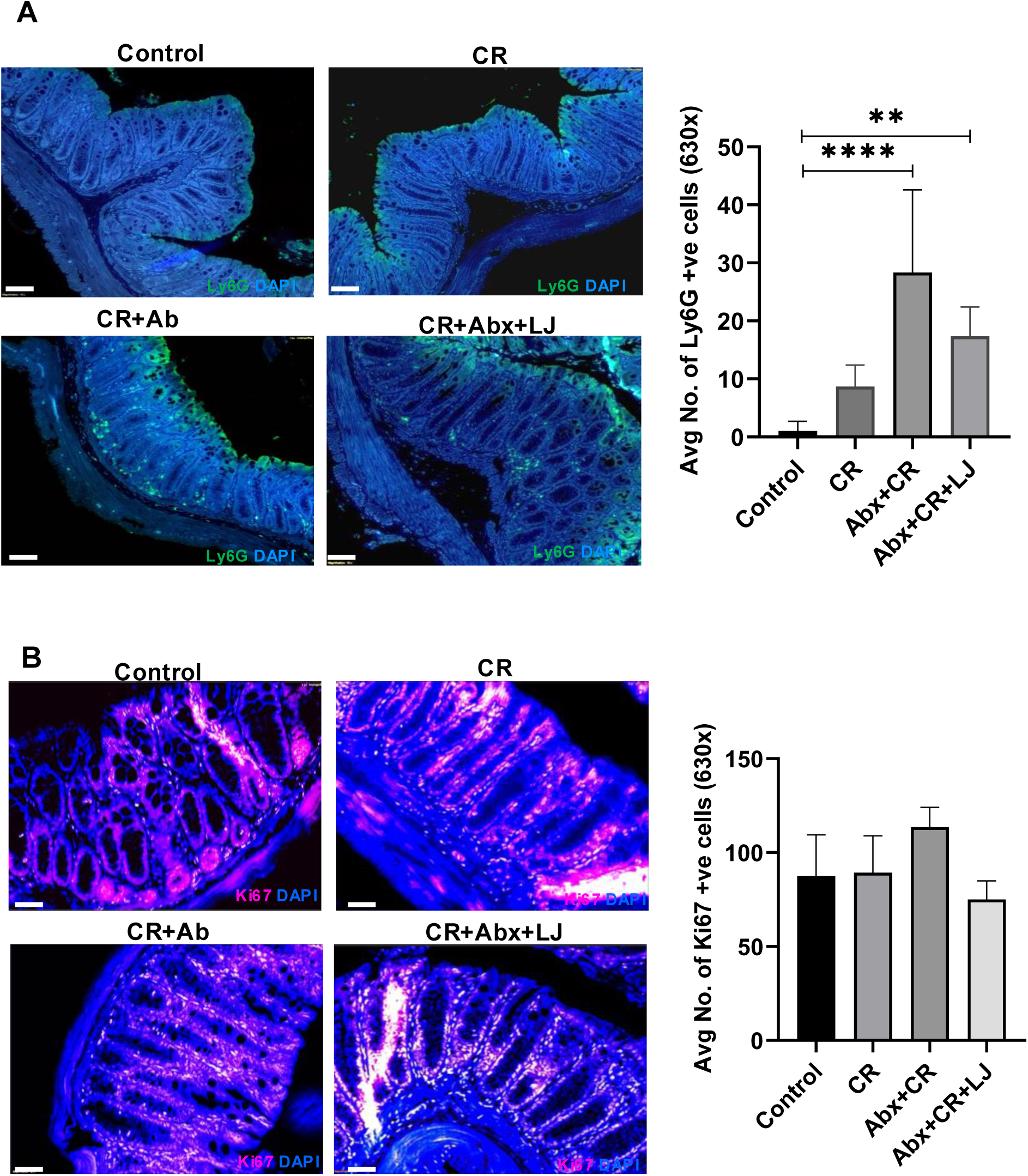
Impact of *L. johnsonii* on neutrophil infiltration and epithelial cell regeneration in colonic tissue. (A) Immunofluorescence staining for Ly6G (neutrophil marker, green) in colonic sections from each treatment group. Nuclei were counterstained with DAPI (blue). Bar graph shows mean ± SEM counts of Ly6G⁺ cells per high-power field (630× magnification) across groups (n = 3 mice/group). Statistical analysis: two-way ANOVA; \*\*\*\**p* < 0.0001, \*\**p* < 0.01. (B) Immunofluorescence staining for Ki67 (proliferation marker, red) in colonic sections from each group, with DAPI (blue) counterstain. Bar graph quantifies Ki67⁺ cells per high-power field (630× magnification). No statistically significant differences were observed between groups.

### Supplementary Table Legends

**S1 Table:** CFU/mL of different bacterial strains at OD600 = 1.

| Bacteria | CFU/ml |
| --- | --- |
| *L. johnsonii* | $5.6 \times 10^7$ |
| *L. plantarum* | $7 \times 10^7$ |
| EPEC | $1.22 \times 10^6$ |
| *C. rodentium* | $1.1 \times 10^8$ |

**S2 Table:** Concentration (µg) of antibiotics used for susceptibility testing.

| Antibiotic | Concentration<br>µg/100µl |
| --- | --- |
| Ampicillin | 10 |
| Kanamycin | 30 |
| Gentamicin | 10 |
| Vancomycin | 30 |

**S3 Table:** Concentration of antibiotic cocktail administered orally (single dose) and through drinking water over 7 days for microbiota depletion.

| **S.No** | **Antibiotic cocktail** | **Orally administered (Single dose)** | **Drinking water (7 days)** |
| --- | --- | --- | --- |
| 1 | Neomycin | 2mg/ml | 1mg/ml |
| 2 | Gentamycin | 2mg/ml | 1mg/ml |
| 3 | Metrodinazole | 2mg/ml | 1mg/ml |
| 4 | Streptomycin | 2mg/ml | 1mg/ml |
| 5 | Vancomycin | 1mg/ml | 0.5mg/ml |

**S4 Table:**
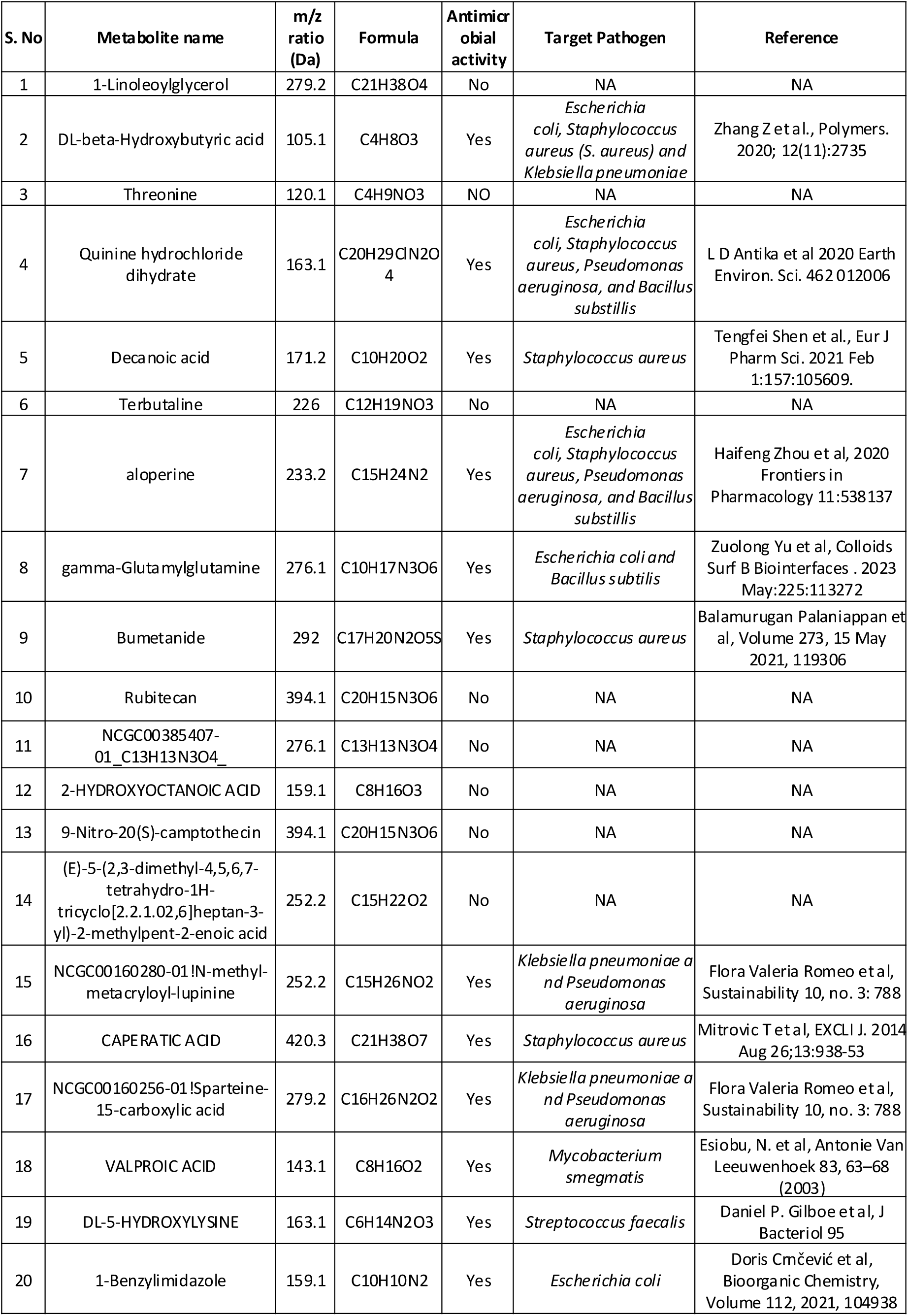
List of major annotated metabolites from the active S6 fraction of L. johnsonii. Eleven compounds exhibited reported antimicrobial activity, including four with activity against E. coli.

